# A Versatile Nanoluciferase Reporter Reveals Structural Properties Associated With a Highly Efficient, N-Terminal *Legionella pneumophila* type IV Secretion Translocation Signal

**DOI:** 10.1101/2022.05.25.493526

**Authors:** Yoon-Suk Kang, James E. Kirby

**Affiliations:** Department of Pathology, Beth Israel Deaconess Medical Center, Boston, MA; Harvard Medical School, Boston, MA

## Abstract

Many gram-negative pathogens rely on type IV secretion systems (T4SS) for infection. One limitation in the field has been the lack of ideal reporters to identify T4SS translocated effectors and study T4SS function. Most existing reporter systems make use of fusions to reporter proteins, for example, β-lactamase, to detect translocated enzymatic activity inside the host cell. However, these systems require costly substrates, complex procedures to separate eukaryotic cytoplasm for analysis, and/or are insensitive. Here, we developed and characterized a novel reporter system using nanoluciferase (NLuc) fusions to address these limitations. Serendipitously, we discovered that Nluc itself is efficiently translocated by *L. pneumophila* T4SS in an IcmSW chaperone-dependent manner via an N-terminal translocation signal. Extensive directed and random mutagenesis in the NLuc N-terminus revealed a critical α-helical domain spanning D5 to V9, as mutations that are predicted to disrupt this α-helix were translocation defective. Notably, NLuc was capable of translocating several proteins examined when fused to the N or C-terminus, while maintaining robust luciferase activity. In particular, it delivered the split GFP11 fragment into J774 macrophages permanently transfected with GFPopt, thereby resulting in *in vivo* assembly of superfolder GFP. This provided a bifunctional assay in which translocation could be assayed in by fluorescence microplate, confocal microscopy, and/or luciferase assay. We further identified an optimal NLuc substrate, which allowed a robust, inexpensive, one-step, high throughput screening assay to identify T4SS translocation substrates and inhibitors. Taken, together NLuc provides both new insight into and tools for studying T4SS biology.

**Importance:** Type IV secretion systems (T4SS) are used by gram-negative pathogens to coopt host cell function. However, the translocation signals recognized by T4SS are not fully explained by primary amino acid sequence, suggesting yet to be defined contributions of secondary and tertiary structure. Here, we unexpectedly identify nanoluciferase (NLuc) as an efficient IcmSW-dependent translocated T4SS substrate and provide extensive mutagenesis data suggesting that the first N-terminal, alpha helix domain is a critical translocation recognition motif. Notably, most existing reporter systems for studying translocated proteins make use of fusions to reporters to permit detection of translocated enzymatic activity inside the host cell. However, existing systems require extremely costly substrates, complex technical procedures to isolate eukaryotic cytoplasm for analysis, and/or are insensitive. Importantly, we find that NLuc provides a powerful, cost-effective new tool to address these limitations and facilitate high throughput exploration of secretion system biology.

## Introduction

Bacterial type IV secretion systems (T4SSs) are macromolecule delivery machines consisting of multi-subunits that span the inner and outer cell membranes of gram-negative pathogens. They are closely related to bacterial conjugation machines (1, 2). A number of gram-negative bacterial pathogens use T4SS to transfer protein and/or DNA into eukaryotic cells, where these effectors then facilitate intracellular or extracellular bacterial replication through modulation of phagosome maturation and/or the immune response (1, 3, 4). T4SS have defining roles in infections caused by pathogens such as *Legionellales, Bartonella* spp., *Brucella* spp., *Bordetella pertussis*, *Helicobacter pylori, Anaplasmataceae*, and *Rhizobium radiobacter* (4–10).

To investigate the translocation of specific bacterial effectors into host cells, a variety of reporter systems have been developed. Fusion of bacterial effectors with adenylate cyclase (CYA) or β-lactamase (TEM-1) reporters have been the two mainstays used in investigation of type III, IV, and VI secretion systems (11–13). In particular, calmodulin-dependent adenylate cyclase has proven sensitive (14, 15) and specific for detecting translocation, the high specificity based on CYA only demonstrating enzymatic activity after combining with calmodulin in the host cell cytoplasm. However, these systems have several liabilities: for CYA, the time-consuming steps needed to extract intracellular cAMP for analysis by immunoassay and, for TEM-1, the high cost of the CCF2/AM fluorescent substrate and the complexity of substrate preloading into host cells.

We therefore set out to develop a next generation reporter system to address these deficiencies. Nanoluciferase (NLuc) is a small 171 amino acid monomer, which was co-evolved from the 19kD OLUC-19 luminescent protein (16) from the deep-sea shrimp, *Oplophorus gracilirostris*, along with a derivatized coelenterazine substrate to optimize luminescence output (17). Notably, it has enhanced structural stability, and provides over 150-fold higher signal than the much larger firefly luciferase widely used in bioimaging and bioreporter studies (18–20). Furthermore, the furimazine substrate optimized for NLuc has low autoluminescence, enhancing the signal to noise ratio for this system. Based on these attributes, we hypothesized that NLuc might prove an optimal translocation reporter for T4SS. Here, we therefore characterize use of NLuc as a T4SS translocation reporter and further investigate a highly efficient translocation signal unexpectedly identified in the N-terminus of NLuc itself.

## Results

### NLuc as a T4SS translocation reporter

To investigate the potential of NLuc as a translocation reporter, we first constructed a fusion with RalF, a well-known T4SS translocated effector from *L. pneumophila* (13). In this constructs, the RalF protein was fused to the C-terminus of NLuc to preserve RalF’s C-terminal translocation signal (13). A A 3X-FLAG epitope was also added to the N-terminus of the fusion protein to facilitate western blot detection.

J774A.1 macrophages were then infected with either the T4SS-competent *L. pneumophila* strain, Lp02, or T4SS-defective strain, Lp03 (*dotA* mutant), expressing the fusion protein. At 6 h post infection, prior to significant bacterial replication and potential phagosomal degradation of T4SS-defective Lp03, translocation was measured in saponin extracts of infected cultures by western blot (Fig. 1A). Of note, saponin, a cholesterol-dependent detergent, is used to liberate selectively eukaryotic cytoplasmic contents without solubilizing bacteria such as *L. pneumophila*, which do not contain cholesterol in their cell membranes. Therefore, in saponin extracts of eukaryotic cells, translocated protein is collected in the extract supernatant, while intact bacteria are pelleted and can be separately assayed as a control for protein expression levels in infecting bacteria.

**Figure 1.**
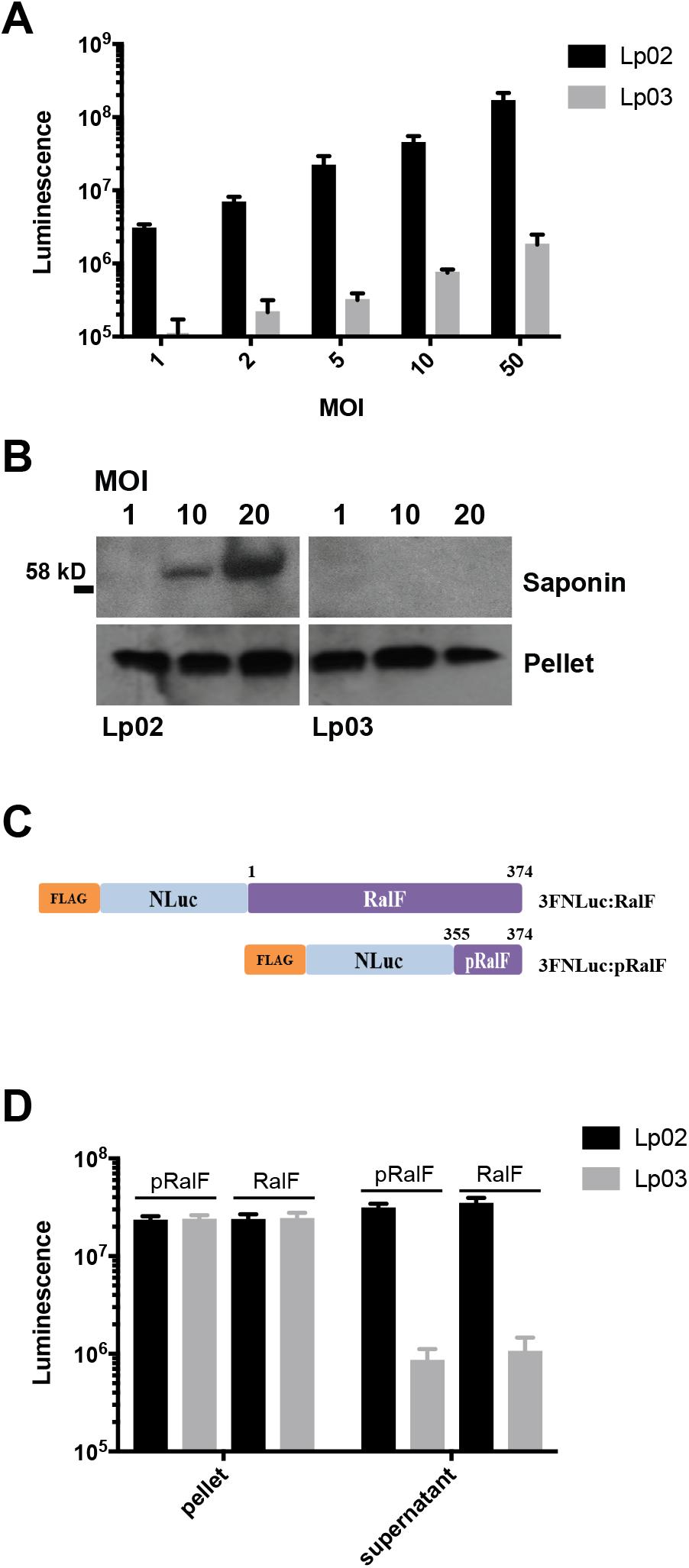
Translocation of nanoluciferase:RalF fusion proteins into J774A.1 macrophages. Macrophages were infected with different multiplicities of infection (MOIs) of wild-type (Lp02) or *dotA* T4SS-incompetent (Lp03) *L. pneumophila* expressing an 3xFlag(3F):NLuc:RalF protein fusion. **(A)** Western blot detection of translocated 3F:NLuc:RalF fusion protein with anti-FLAG tag antibody 6 h post infection. Translocated protein is found in the extracted eukaryotic cytoplasm (supernatant), while bacterial-associated, untranslocated protein remains in the bacterial pellet. **(B)** Detection of translocated luciferase activity under similar conditions. **(C)** Diagram of NLuc constructs fused with whole RalF (amino acids 1-374) or partial RalF (C-terminal amino acids, 355-374), designated NLuc:RalF or NLuc:pRalF, respectively. **(D)** Translocation of NLuc:RalF and NLuc:pRalF expressed in Lp02 or Lp03 following a 6 h infection at an MOI of 10. Mean luminescence and standard deviation from three independent experiments are shown.

We found that FLAG-tagged protein was only detected in eukaryotic cytoplasmic extracts during Lp02 infection, but not during Lp03 infection, in rough proportion to multiplicity of infection, supporting translocation of the NLuc:RalF fusion protein (Fig. 1A). Importantly, similar levels of FLAG epitope signal were detected in extract pellets from Lp02 and Lp03, indicating that differences in protein expression in these strains could not account for pronounced differences in protein detected in saponin extracts. Taken together, these results indicated that the NLuc protein could be transported by *L. pneumophila* T4SS when fused to RalF.

We then examined whether NLuc would retain its luciferase activity in a translocated fusion protein. Indeed, in eukaryotic cytoplasmic extracts, a robust, T4SS-dependent luminescence signal was observed on addition of furimazine substrate (Fig. 1B). Signal was approximately 100-fold higher in the wild type than *dotA* T4SS mutant background and correlated with multiplicity of infection (Fig. 1B). In contrast, similar levels of NLuc activity were observed in extract pellets from Lp02 and Lp03 infection. These results suggested that the NLuc fusion protein was translocated into the macrophage cytoplasm in a manner that also preserved its luciferase activity.

### NLuc contains an intrinsic T4SS translocation signal

Previously, the twenty amino acid C-terminus of *L. pneumophila*, RalF effector was found to be both necessary and sufficient for translocation (13). However, translocation mediated by this C-terminus alone occurred with reduced efficiency. Based on these prior observations, we tested whether the C-terminus of RalF was similarly necessary and sufficient for NLuc translocation (Fig. 1C,D). Unexpectedly, essentially identical T4SS-dependent NLuc signal was observed in eukaryotic cytoplasmic extracts 6 h post infection with *L. pneumophila* expressing NLuc fused with either the C-terminal 20 amino acids of RalF (NLuc:pRalF) or full length RalF (NLuc:RalF). This result suggested that NLuc either compensates for the lower translocation efficiency previously associated with the C-terminus of RalF or alternatively provides its own translocation signal.

To investigate the possibility of an intrinsic translocation signal in NLuc, we expressed NLuc with either an N- or C-terminal 3X FLAG tag (Fig. 2A). At 6 hpi, we detected essentially identical T4SS-dependent translocation of these constructs by western blot (Fig. 2B). Similar results were observed for luciferase activity (Fig. S1A). NLuc was translocated just as efficiently in the absence of an N- or C-terminal 3X FLAG epitope based on nanoluciferase assay, suggested that the 3X FLAG itself did not contribute to the translocation phenotype (Fig. S1A). Taken together, these results suggested the presence of an intrinsic T4SS translocation signal in NLuc, and furthermore that the 3X-FLAG tag interfered with neither translocation nor luciferase activity, irrespective of location.

**Figure 2.**
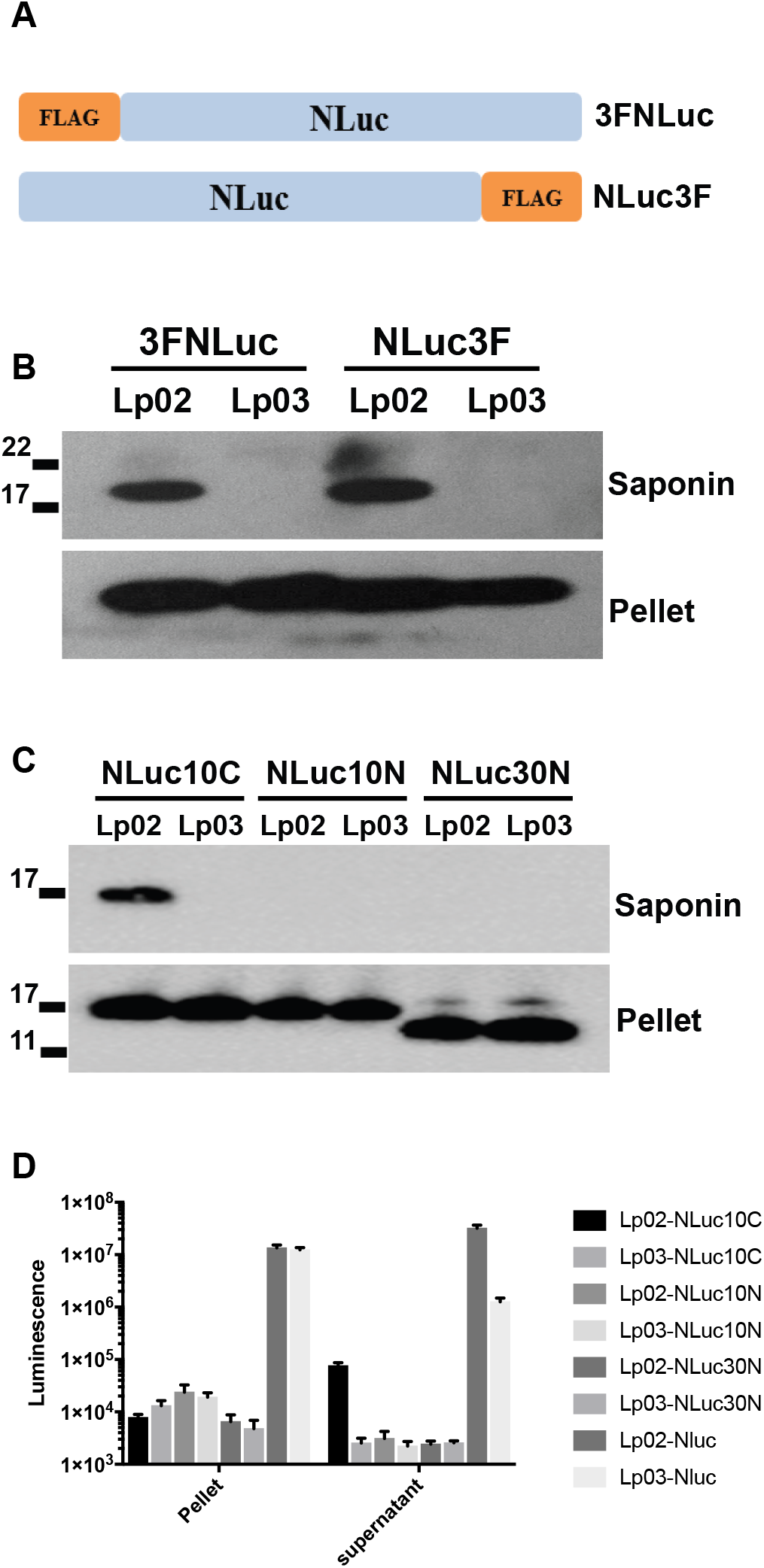
Intrinsic translocation signal in NLuc. **(A)** Diagram of NLuc constructs containing a 3xFLAG tag in the N-terminus (3FNLuc) or C-terminus (NLuc3F). **(B)** Western blot detection of translocation of 3FNLuc and NLuc3F expressed by T4SS-competent (Lp02) and T4SS-incompetent (Lp03) 6 h post infection detected using anti-FLAG antibody. **(C)** Translocation of NLuc10C (deleting the 10 C-terminal amino acids), NLuc10N (deleting the 10 N-terminal amino acids) or NLuc30N (deleting the 30 N-terminal amino acids) expressed by Lp02 or Lp03 6 h post infection detected using anti-FLAG antibody (constructs as diagrammed in Fig. S1B). **(D)** Detection of translocated luciferase signal in extract supernatant and pellets from J774A.1 cells infected for 6 h with Lp02 and Lp03 strains expressing C- and N-terminal NLuc truncations. Mean and standard deviations from three independent experiments are shown.

To identify the T4SS translocation signal in NLuc, we created a series of C- and N-terminal deletions of 3X-Flag-NLuc (Fig. S1B) and assayed for translocation 6 h post infection. Deletion of the 10 C-terminal amino acids was associated with modest T4SS-dependent protein translocation, as assessed by western blot (Fig. 2C) and luciferase activity (Fig. 2D). In contrast, deletion of either the 10 or 30 N-terminal amino acids abolished translocation. These data are consistent with a T4SS translocation signal residing in the N-terminus of NLuc. Notably, although protein expression appeared relatively unaffected as assessed by western blot, the luciferase activity observed in bacterial pellets of all deletion constructs was substantially reduced, suggesting that both the N and C termini of NLuc are required for efficient luciferase activity.

### Characterization of the N-terminal translocation signal

We next asked whether the N-terminal 30 amino acids of NLuc (partial NLuc or pNLuc) were necessary and sufficient for T4SS-dependent translocation. To do so, we constructed fusions of TetR and TEV protease proteins fused to pNLuc with a N- or C-terminal 3X-FLAG tag (Fig. S2A). Importantly, neither TetR nor TEV protease tagged with an N-terminal 3X-FLAG alone were translocated by themselves (Fig. S2B). However, both proteins were translocated in a T4SS-dependent manner when fused with N- or C-terminal pNLuc (Fig. S2C, D). Interestingly, pNLuc:TetR was translocated more efficiently than TetR:pNLuc, while TEV:pNLuc was translocated slightly more efficiently than the pNLuc:TEV. These results suggested that the specific fusion protein partner and its location relative to NLuc can influence either the availability of the translocation signal and/or efficiency of translocation.

### Dependence of NLuc translocation on IcmSW components of the type IV coupling transport complex

The chaperones, IcmS, IcmW, and LvgA, facilitate translocation of *L. pneumophila* effectors that lack canonical C-terminal E-block motifs (described further below) through participating in substrate recognition in the type IV coupling transport complex (21–23). We therefore examined their contribution to NLuc translocation through use of genetic knockouts. Interestingly, translocation was partially reduced in both isogenic *icmW* and *icmS*, but not in the *lvgA* mutant strains (Fig. 3A). In the *icmSW* double knockout, translocation efficiency was reduced by ∼84%, a defect that was fully complemented by an *icmSW* expression plasmid. The magnitude of *icmSW* translocation dependence is consistent with previous observations for translocation substrates that specifically bind to the IcmSW chaperone complex (24, 25).

**Figure 3.**
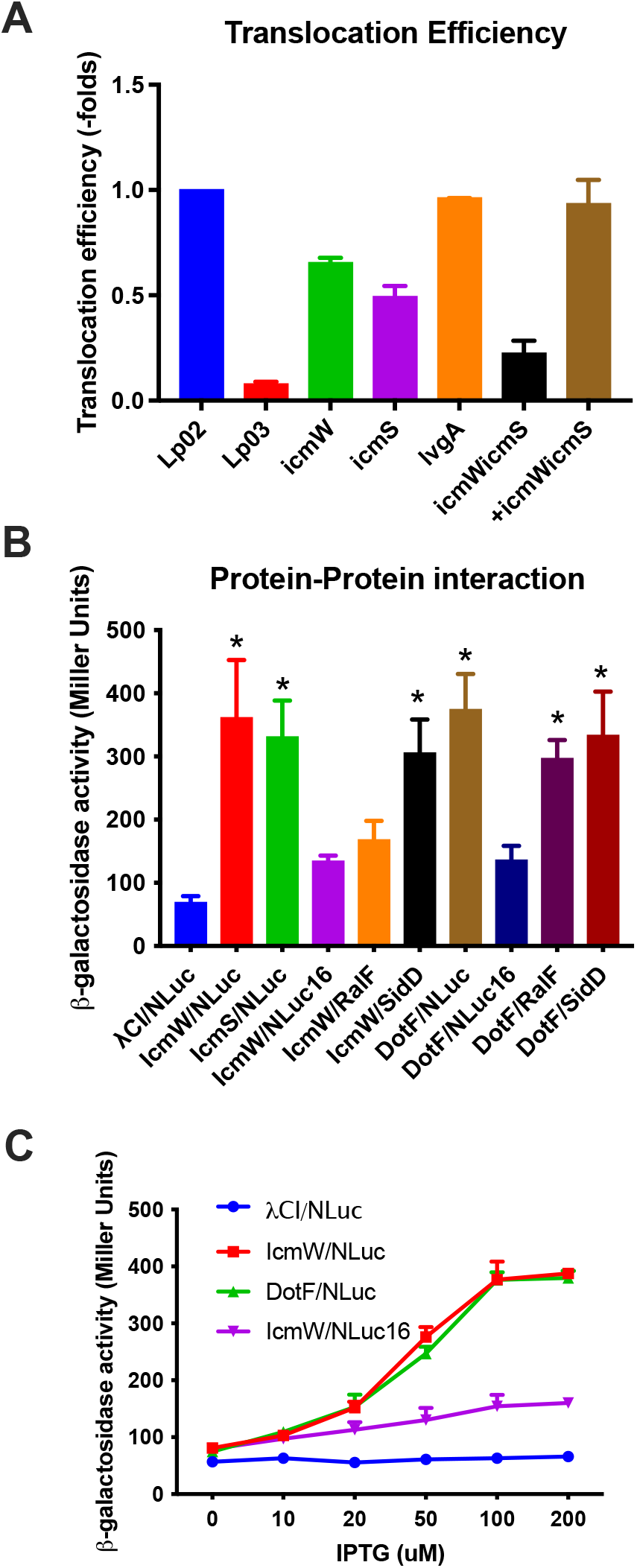
Dependence of NLuc Translocation Signal on IcmSW. **(A)** Luminescence-based translocation efficiency of NLuc in *L. pneumophila* Lp02, Lp03 *dotA*, Δ*icmW*, Δ*icmS*, Δ*lvgA*, Δ*icmSW*, and Δ*icmSW* complemented with plasmid expressed *icmSW*. Extract supernatants and pellets were obtained 6 h post infection at an MOI of 10. Translocation efficiency in mutants was normalized to signal from Lp02. Shown are the mean and standard deviation from two independent experiments. **(B)** Bacterial two-hybrid (BACTH) assays to detect interactions of NLuc, RalF and SidD prey with IcmW, IcmS and DotF bait. Shown are mean and standard deviation of β-galactosidase activity from three independent experiments following induction of bait and prey fusion constructs with 100 µM IPTG. Statistical significance was determined using one way ANOVA test with Holm-Sidak’s multi-comparison post hoc test, * indicates adjusted *P* < 0.05 for the comparison of the indicated condition with the λCI:β-flap/Nluc:αNTD expression vector combination (abbreviated λCI/NLluc) negative control. **(C)** Interaction of NLuc or NLuc16 with IcmW, DotF, or β-flap negative control. The mean and standard deviation of β-galactosidase activity for two independent experiments are plotted versus IPTG concentration.

Therefore, we next considered whether the NLuc and its N-terminus might directly interact with IcmSW and other proteins components of the T4SS apparatus, as described for other effectors. To do so, we made use of a previously described bacterial two-hybrid system to test for such protein-protein interaction (26, 27). In this system, interaction of one potential protein partner, fused to λCI, with a second protein, fused to the N-terminal domain (NTD) of the of α subunit of RNA polymerase, leads to transactivation of a β-galactosidase reporter. Using this method, we detected strong interaction between NLuc:αNTD and λCI:IcmW, and between NLuc:αNTD and λCI:DotF (Fig 3B). Readout was dependent on IPTG concentrations used for induction of protein partners, supporting specificity of findings (Fig. 3C). Importantly, a negative control for baseline expression was established by testing co-expression of presumptively non-interacting λCI fused to β-flap region of the RNA polymerase β-subunit and Nluc:αNTD (Fig 3B, λCI/NLuc), resulting ia comparatively low baseline transactivation that was not similarly induced by IPTG (Fig. 3C). As positive controls, we also examined interactions described previously (25, 28–30), detecting strong interaction (transactivation) between RalF and DotF; SidD and DotF; and SidD and IcmW. In contrast, as expected, we only detected weak, not-statistically significant interaction between the IcmSW-independent effector, RalF, and IcmW (Fig. 3B).

Importantly, when testing fusions with a version of NLuc with a N-terminal 15 amino acid truncation (NLuc16:αNTD), we only observed weak, non-significant interaction with λCI:IcmW and λCI:dotF (Fig. 3B, C). These results suggested that the N-terminal translocation domain of NLuc contributed to interactions with the T4SS machinery components examined.

### Definition of translocation signal through random mutagenesis and targeted interruption of predicted secondary structure

Based on the preceding results, we examined the 20 N-terminal amino acids for motifs previously associated with translocation. Previous studies identified a so-called E-block within C-terminal T4SSb-dependent translocation signals in *L. pneumophila* (13, 31–33). This negatively charged motif, defined by enrichment in glutamate and aspartate residues (33), has been proposed to interact with cytoplasmic facing, positively-charged amino acids of the DotM, T4SS protein, to initiate translocation (21).

The presence of aspartic and glutamic acid residues in the N-terminus of NLuc (Fig. 4A, S3A), suggested potentially similar functionality. To characterize their contribution to translocation, we replaced these negatively charged amino acids sequentially with charge-neutral, non-polar alanine and/or glycine residues (Fig. 4A). At 6 hpi, T4SS-dependent translocation of these NLuc mutants was measured (Fig. 4B) by luminescence signal and was noted to be reduced in a modest, step-wise fashion from ∼28% to ∼44% with single (E6G or D7A), double (E6G/D7A or E6G/D11A), or triple mutations (E6G, D7A, and D11A). Importantly, intrinsic luciferase activity of NLuc mutants was no different from wild type NLuc (Fig. 4C, bacterial pellets), suggesting lower luciferase activity in eukaryotic cytoplasmic extracts resulted from reduced translocation. Based on a partial phenotype, we infer that other features in the N-terminus of NLuc must also contribute significantly to its translocation.

**Figure 4.**
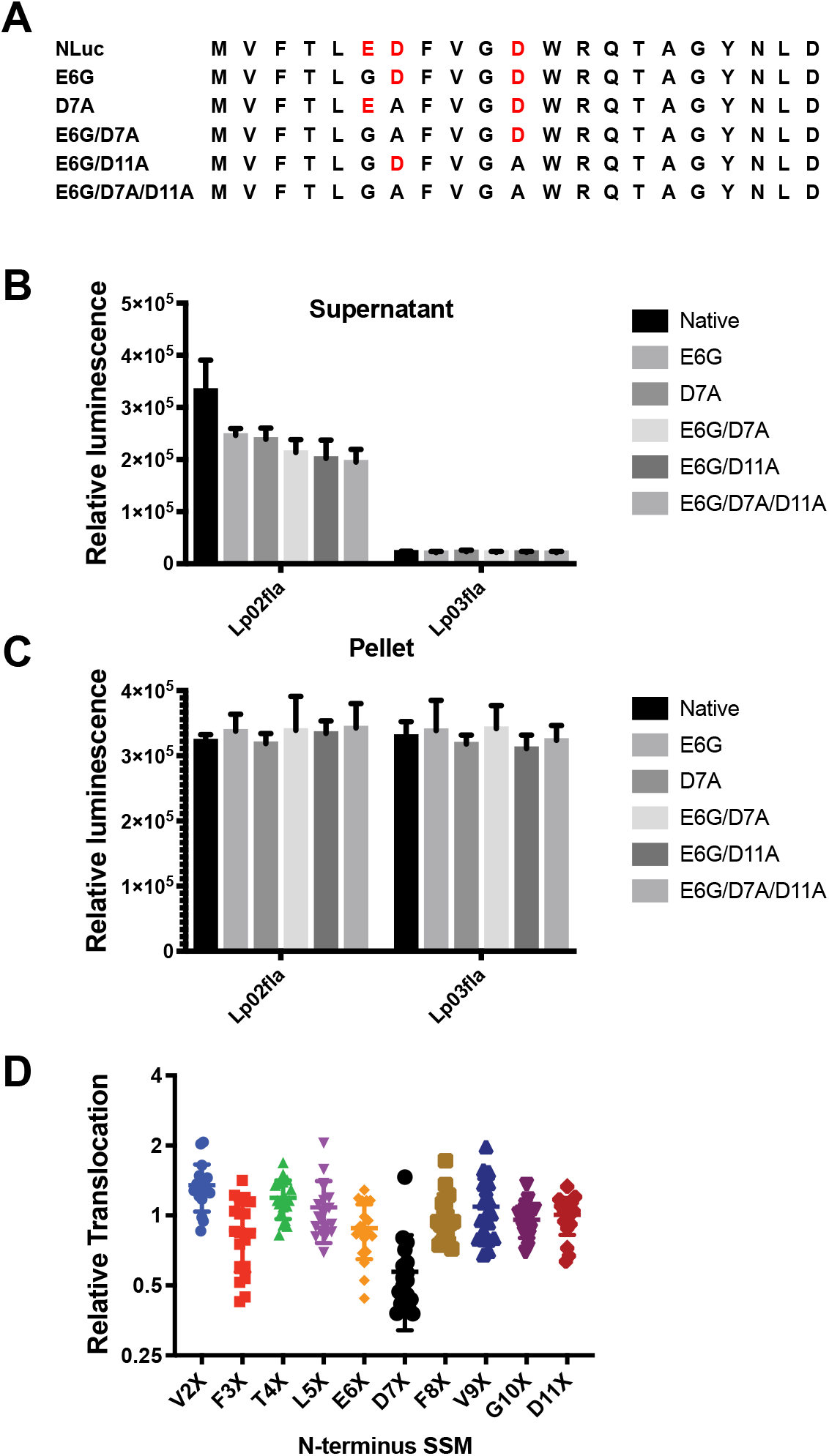
N-terminal translocation signal does not substantially depend either the presence of negatively charged amino acids or its primary amino acid sequence. **(A)** Negatively charged amino acids (red type) in the N-terminus of NLuc were mutated to neutral alanine or glycine. J774A.1 cells were infected for 6 h at an MOI of 10 with *L. pneumophila* strains expressing NLuc with indicated mutations. Saponin extract supernatant **(B)** and pellet **(C)** were then assayed for nanoluciferase activity. Data shown are the mean and standard deviation of three independent experiments. **(D)** Scanning, random mutagenesis of V2-D11 amino acids in NLuc. Relative translocation of each mutant was calculated based on the ratio of luciferase activity in the saponin extract to bacterial pellet 6 h post infection with Lp02 expressing the indicated mutant at an MOI of 10. Values were normalized to translocation of wild type NLuc and are the average translocation values from two independent experiments.

To investigate more broadly the potential involvement of specific amino acid sequences in the N-terminal translocation signal, we replaced each amino acid from V2 to D11 sequentially through randomization of each respective codon. We then compared the translocation efficiencies normalized to expression in bacterial pellets (to control for effects of mutations on intrinsic luciferase activity) during J774A.1 cell infection. Surprisingly, all mutations were well tolerated except positions 3F, 6E, and 7D where reduced translocation of up to 70% was observed (Fig. 4D, Table S1). However, no single substitution was able to fully block translocation. Mutations either variable decreased, had no effect, or in some cases, increased absolute luminescence (Table S1).

Due to only partial abrogation of translocation by single amino acid mutations, we considered whether secondary structure might contribute to NLuc recognition by T4SS. The RaptorX convolutional neural network-based, secondary-structure prediction program (34) predicted an α-helical region in the N-terminus of NLuc corresponding to the surface-exposed α-helix observed in the crystal structure of NLuc in PDB 5IBO (Table S2) (35). Single amino acid substitutions such as E6G, D7A, D11A or their combinations did not disrupt the predicted α-helical structure. However, Δ6E, Δ7D, and Δ6E7D deletions did (Table S2) and experimentally were correspondingly noted to decrease normalized translocation up to 94% for Δ7D, with subtraction of background signal from the Lp03 T4SS incompetent control (Fig. 5A). A similar elimination of α-helical structure was predicted by AlphaFold2 (Fig. S3) (36). Interestingly, all three deletions increased luminescence signal roughly four to six-fold in pellets (Table S1) suggesting opposite effects on enzymatic and translocation activities.

**Fig. 5.**
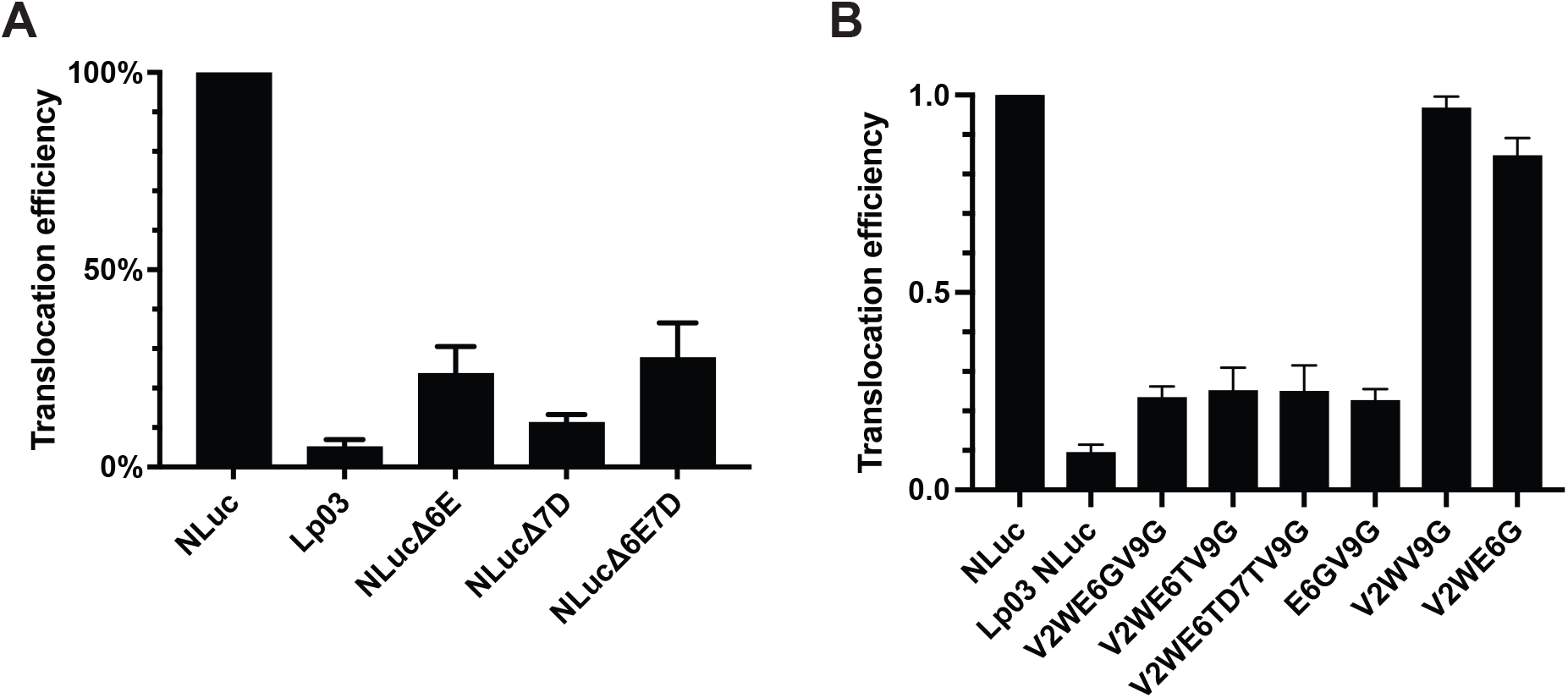
Predicted disruption of the α-helical region in the N-terminus of Nluc abrogates translocation. **(A)** Translocation efficiency of parental NLuc in Lp02 or Lp03 and NLuc1Δ6E, NLuc1Δ7D, and NLuc1Δ6E1Δ7D expressed in Lp02. Translocation efficiencies of mutants are the ratios of luciferase activity in the saponin extract supernatant to bacterial pellet normalized to the translocation efficiency of wild type NLuc, testing performed 6 h post infection of J774A.1 cells at an MOI of 10. Bars are the mean and standard deviation from two independent experiments. **(B)** Translocation efficiency of wild type NLuc and the multi-site mutants indicated, expressed in Lp02. Bars shown are the mean and standard deviation from three independent experiments.

Based on these data, we designed degenerate primers to randomly mutagenize positions V2 through W12. Over 86% of the ∼7000 clones analyzed showed >10-fold decreased luminescence and were not further analyzed (Fig. S4). We selected ∼30 NLuc mutants showing high luciferase activity and evaluated translocation efficiency (Table S2). Interestingly, these highly active luciferase variants showed changes in only one to four amino acid positions compared with the native nanoluciferase sequence, suggesting relative stringent conservation requirements in this region for optimal luciferase activity. Based on observed mutations, we also constructed V2WE6G and E6GV9G. Among all variants examined, V2WE6TV9G; V2WE6TD7TV9G; V2WE6GV9G, and E6GV9G showed ∼5-fold decreased translocation compared to parental NLuc (Fig. 5B) However, translocation of V2WV9G and V2WE6G were unaffected (0.97 translocation efficiency). Notably, all of the former, and V2WV9G, showed a predicted disruption of α-helical structure based on RaptorX, while none showed disruption based on AlphaFold2 (Fig. 5B). Taken together our results suggested that the surface-exposed, N-terminal α-helical secondary structure may be a critical component of the *icmSW*-dependent T4SS signal.

### Developing a fluorescent T4SS translocation reporter using split, superfolder GFP

Based on its native T4SS signal, we next considered whether it would be possible to tether the Nluc protein to a split-GFP superfolder (sfGFP) fluorescent reporter (37) to provide a specific, and temporal readout of T4SS translocation without the need for extraction of eukaryotic cytoplasm. Utility of split-sfGFP is predicated on *in vivo* self-assembly of the individually non-fluorescent, 16 amino acid GFP11 fragment and complementary GFP1-10 OPT (GFPopt) domains with resulting restoration of GFP fluorescence (37, 38). We constructed a split-sfGFP reporter system on this basis.

We initially checked *in vivo* assembly of GFP11 fused with either pRalF or NLuc (GFP11:pRalF and NLuc:GFP11, respectively) and GFPopt in *E. coli* (Fig. S5A). Co-expression of either of the two GFP11 fusions with GFPopt resulted in increased fluorescent signal (*P* < 0.05), i.e., significantly above the background signal represented by expression of GFPopt alone. These results indicated *in vivo* reconstitution of functional sfGFP in *E. coli* when GFP11 was available in the form a fusion protein with either pRalF or NLuc translocation signals.

Based on this data, J774A.1 host cells expressing GFPopt were infected with *L. pneumophila* strains expressing either NLuc:GFP11 or GFP11:pRalF in 96-well microplate assays (Fig. 6A). At 24 h post infection, we observed a significant (Fig. 6B, *P* < 0.001) T4SS-dependent GFP signal, i.e., for Lp02, but not for Lp03, expressing either NLuc:GFP11 or GFP11:pRalF fusion proteins. As expected, control strains expressing GFP11 in the absence of a translocation signal did not show Lp02-dependent signal. Of interest, signal for Lp02 expressing NLuc:GFP11 was 40% greater signal than Lp02 expressing GFP11:pRalF, suggesting that NLuc provided a stronger translocation signal for the GFP11 reporter fragment. Comparisons between signal from Lp02 and Lp03 infection yielded a Z’ of 0.32±0.01 for Nluc:GFP11 and -0.11±0.28 for GFP11:pRalF, respectively (Fig. S5B), suggesting that the former, but not the latter, had robustness compatible with use in high throughput translocation inhibition assays. Presumably, there could be additional improvement in Z’ with further assay optimization.

**Figure 6.**
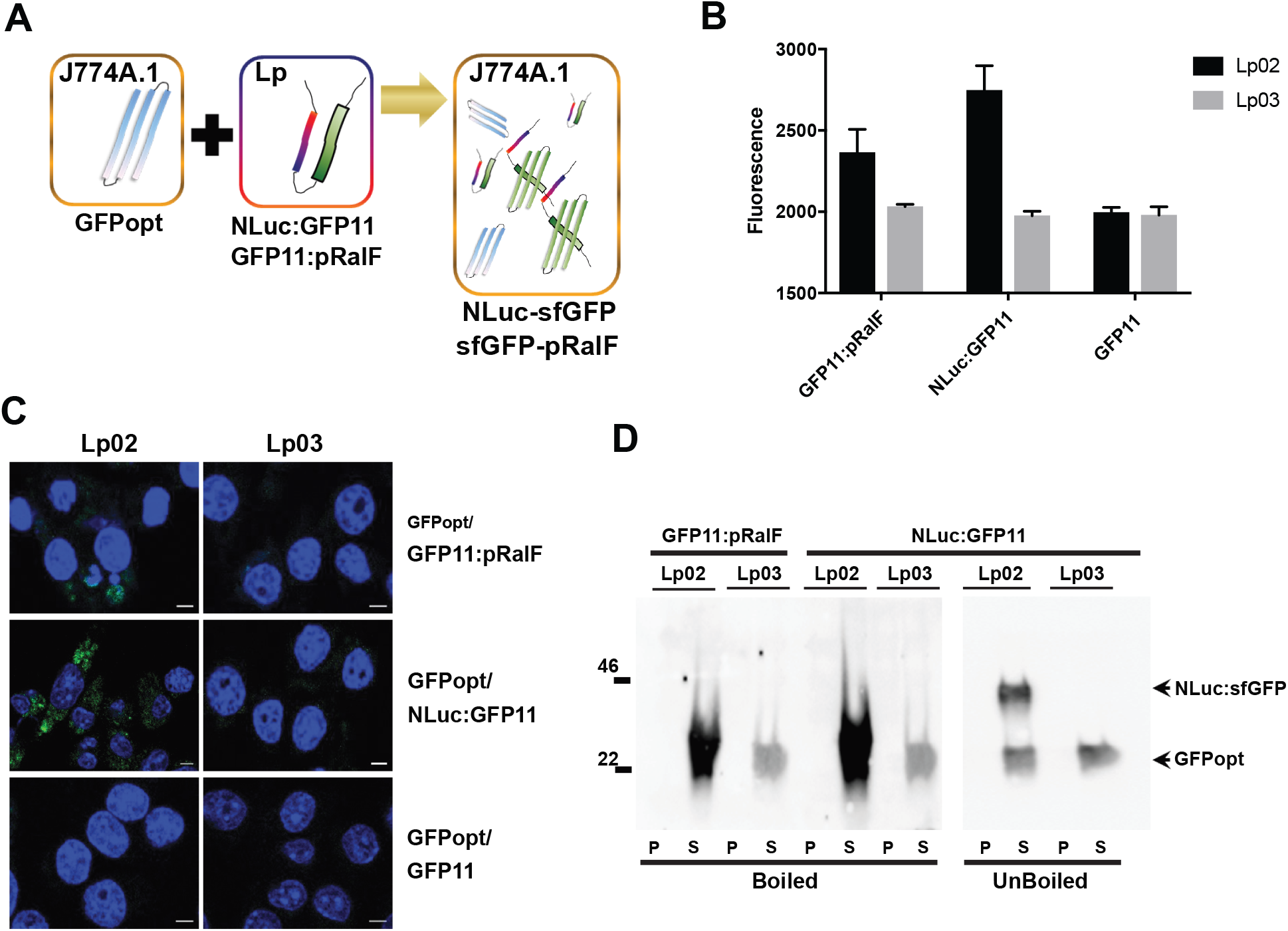
Combining superfolder GFP and NLuc systems to build a bifunctional, internally-orthogonal T4SS-dependent high throughput translocation assay. **(A)** Non-fluorescent GFPopt is expressed constitutively in J774A.1 macrophages. NLuc:GFP11 or GFP11:pRalF translocated into infected macrophages from *L*. *pneumophila* combines with GFPopt to form fluorescent sfGFP detectable in microplate assay or by confocal microscopy. Translocated luciferase activity can be detected separately as an orthogonal measure after extraction of eukaryotic cytoplasm or in a one-step assay described in Fig. 7. **(B)** In the experiment shown, J774A.1 GFPopt macrophages, seeded in 96-well plates at 1× 10^5^ cells per well, were infected for 24 h with *L. pneumophila* expressing either GFP11:pRalF or NLuc:GFP11. Assembly of sfGFP was quantified by fluorescence. The mean and standard deviation of data from three independent experiments each performed in triplicate is shown. **(C)** Alternatively, assembly of fluorescent sfGFP (green) was detected by confocal microscopy with DAPI counterstain (blue). Scale bars are 5 μm. **(D)** Cytoplasmic extracts and pellets of J774A.1 GFPopt cells infected for 24 h with Lp02 or Lp03 expressing GFP11:pRalF or Nluc:GFP11 were analyzed by western blot using polyclonal anti-GFP antibody. Samples from Nluc:GFP11 infections were loaded onto SDS-PAGE gels in standard SDS loading buffer with (boiled) or without (unboiled) prior incubation at 95°C for 5 min. The ∼46 kDa band observed in the unboiled Lp02 supernatant sample is the size predicted for an assembled complex of Nluc:GFP11 (22 kDa) and GFPopt (24 kDa).

To investigate the utility of these reporters in image-based analysis, we examined infections with these constructs using confocal microscopy (Fig. 6C). Confocal images showed host cell cytoplasmic fluorescent signal during infection of J774A GFPopt cells with Lp02 but not with Lp03 expressing either GFP11:pRalF or NLuc:GFP11 fragments. However, signal was not observed when the GFP11 fragment was expressed alone. This latter control, also shown in Fig. 6B, indicated that GFP signal did not result from leakage of GFP11 into the host cytoplasm as a result of bacterial lysis during the course of infection. Similar to microwell experiments, T4SS-dependent translocation signal was noticeably greater for NLuc::GFP11 than for GFP11::pRalF.

Western blot analysis provided additional indirect evidence for translocation-dependent assembly of sfGFP in the host cell cytoplasm (Fig. 6D). Notably, during infection with Lp02 expressing GFP11:pRalF, a very strong ∼24kDa band, the expected size of host cell-expressed GFPopt, was detected with polyclonal anti-GFP antibody by western blood after denaturing electrophoresis. In contrast, during Lp03 infection, the 24kDa band was significantly weaker, suggesting much lower amounts of GFPopt protein. We therefore reasoned that T4SS-dependent translocation of GFP11:pRalF led to stabilization and higher accumulation of the 24kDa GFPopt in the context of an assembled sfGFP complex, consistent with fluorescence microscopy results. Presumably, during denaturing conditions used for SDS-PAGE analysis, the larger sfGFP complex was dissociated, and the polyclonal anti-GFP antibody was also unable to detect the very small GFP11 portion of the translocated fusion protein.

Similar T4SS-dependent stabilization of the 24 kDa GFPopt band was observed during infection with *L. pneumophila* expressing NLuc:GFP11. However, if extracts were not boiled prior to gel loading, a strong ∼46 kDa band was now detected during infection with T4SS-compentent Lp02. This band is likely accounted for by formation of the complex between 22kDa NLuc:GFP11 and 24kDa GFPopt. Under these conditions, we infer that remaining weak 24kDa band in extract supernatants observed during both Lp02 and Lp03 infection represents GFPopt protein that did not participate in sfGFP complex formation.

### Single-step, NLuc-based, high throughput screening assay

Furimazine, a coelenterazine analogue, has been the substrate of choice for nanoluciferase-based applications (17–19). However, the high permeability of *Legionella* to furimazine necessitated time-consuming steps to separate eukaryotic cytoplasm from bacteria prior to nanoluciferase-based translocation assessment. We therefore considered whether identification of a less bacterial-penetrant, coelenterazine substrate might enable a simpler, one-step translocation assay.

The bacterial membrane permeability index of available coelenterazine analogs (Table S3) was empirically determined based on a ratio of luminescence from bacteria suspended in PBS to luminescence of bacteria permeabilized in radioimmuneprecipitation (RIPA) buffer containing 1% NP-40 and 1% sodium deoxycholate. Results were normalized to activity of purified nanoluciferase added to PBS and RIPA alone, as RIPA buffer variably inhibited or stimulated activity of nanoluciferase substrates up to 10-fold. Coelenterazine h, i, cp, and n showed the lowest permeability index (three to five-fold lower than furimazine) among nanoluciferase substrates with high signal.

Based on these data, we then compared the ability of high penetrant analogs (coelenterazine f and furimazine) and low penetrant analogs to serve as discriminatory substrates in one-step, NLuc translocation assays in 96-well microplate format. In these one-step assays, substrate was added directly to infected cells at 6 hpi in the presence of 0.2% saponin. Those with low bacterial permeability (h, cp, i, and n) displayed distinct and statistically significant *(P <* 0.0001, Mann-Whitney U Test) T4SS-dependent luminescent signal (Fig. 7A, Table S2). Coelenterazine h was then compared with furimazine tested in a one-step, 96-well high throughput assay format, the former yielding a discriminatory Z’ of 0.52±0.05 and the latter a non-discriminatory Z’ of -5.9±0.3 during comparisons of Lp02 T4SS-competent and Lp03 T4SS-incompetent infection (Fig. 7B). Notably, NLuc:GFP11 performed similarly to NLuc in single step nanoluciferase translocation assays with coelenterazine h substrate (Fig. S5C). This suggests the potential use of bifunctional NLuc:GFP11 in combination with GFPopt J774A.1 cells to provide orthogonal confirmatory readouts for T4SS translocation in the same high throughput screening assay, providing a highly efficient system for studying biochemical or genetic perturbation of T4SS function.

**Figure 7.**
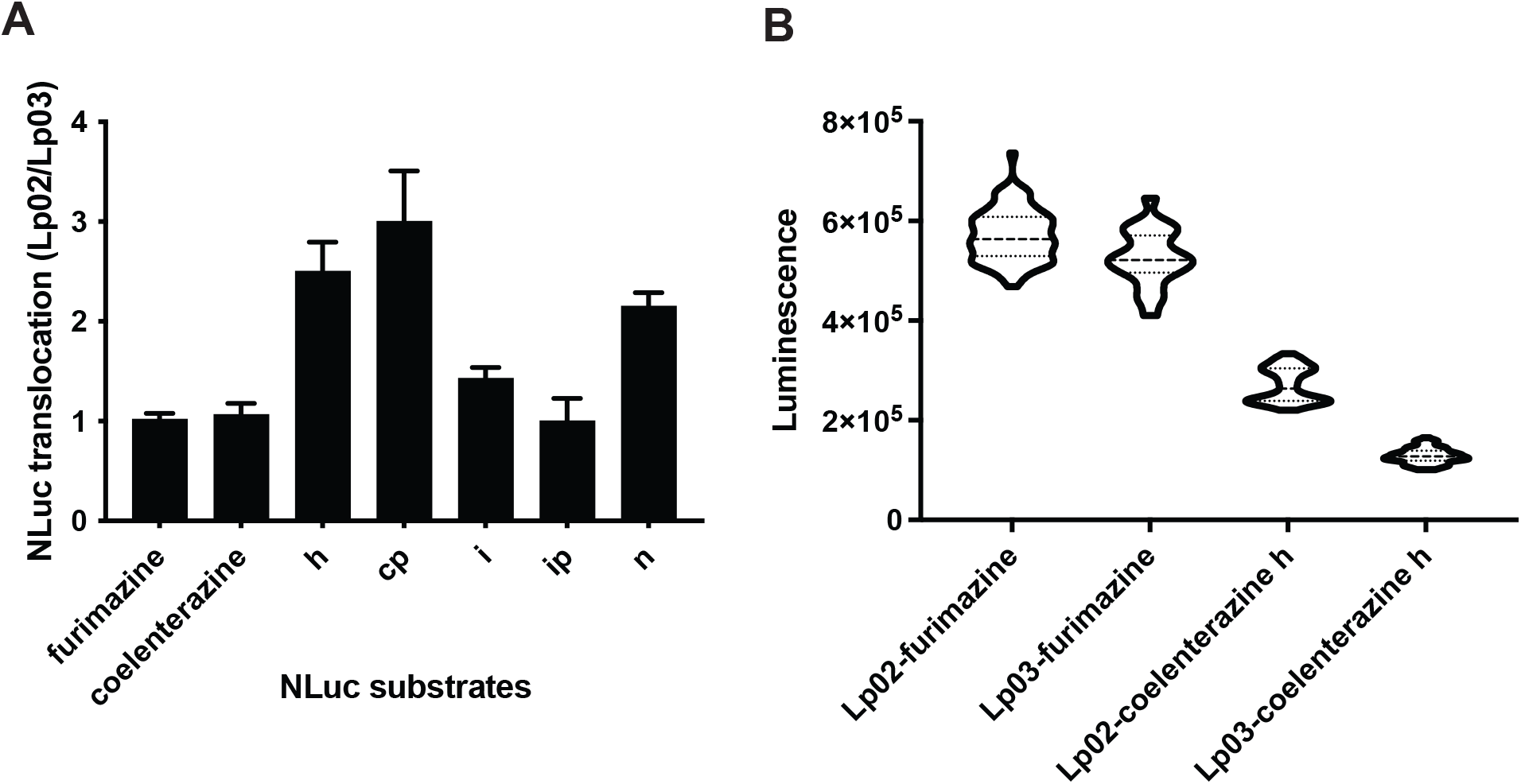
Identification of coelenterazine analogs compatible with a one-step NLuc translocation assay. **(A)** Ratio of luminescence in assay wells after a 6 h infection of macrophages with Lp02 versus Lp03 expressing NLuc on addition of indicated coelenterazine analogues in saponin lysis buffer, i.e., procedurally a single-step translocation assay. Data show are mean and standard deviations for ten determinations obtained during three independent experiments normalized to the furimazine control. **(B)** Luminescence from one-step translocation assays in high throughput format during infection with Lp02 or Lp03 expressing NLuc in the presence of furimazine or coelenterazine h substrates. Violin plots show combined results from two independent experiments, each with 40 technical replicates per variable, which were used to determine the Z’ described in the text. In one-step assays, clear separation between T4SS-competent (Lp02) and -incompetent (Lp03) signal was observed with the coelenterazine h substrate.

## Discussion

Our results identify nanoluciferase (NLuc) as a new reporter for analysis of T4SS-dependent translocation with several propitious properties. The small 17kDa NLuc reporter provided robust translocation signal when fused to both N- and C-termini of proteins examined. It could be detected using a number of inexpensive nanoluciferase substrates or by western blot using epitope tags or nanoluciferase antibody and was compatible with high throughput assay formats. It offered considerable advantage over other systems in current use which require complex techniques for isolation or detection of reporter enzyme substrates at considerable expense.

As NLuc is an evolved, eukaryotic, deep-sea shrimp luciferase, a surprising finding was the presence of a very strong N-terminal, T4SSa-dependent translocation in Nluc itself. Notably, the specific mutations introduced in development of NLuc (16, 17) –– T2V, A6E, R12Q (NLuc numbering) – from *Oplophorus* luciferase coincidentally appeared to enhance translocation efficiency based on our mutagenesis analysis (Table S1) and may have serendipitously contributed to the NLuc translocation signal.

The existence of an N-terminal T4SSb translocation signal in *L. pneumophila* to the best of our knowledge has not been previously established through deletion or mutagenesis experiments. However, of interest, 8 Leg proteins and LepA were previously found to be translocated either as fusions to the N-terminus of CYA or to the C-terminus of TEM β-lactamase (39–41). These results could be consistent with a C-terminal signal that remains active despite downstream fusion protein sequence (40, 41); or potentially, a signal emanating from other regions of these effectors, including the N-terminus. Of note, many studies seeking to identify T4SSa translocated effectors have been based on screens in which candidate effector proteins were fused to the C-terminus of reporters (42, 43), thereby potentially ignoring strong N-terminal signals. Internal recognition sites are also postulated for some IcmSW substrates (25), suggesting C-terminal signals are not obligatory. Therefore, the discovery of a strong N-terminal signal is not completely surprising.

Biological investigation suggested that NLuc translocation was IcmSW-dependent, as ∼90% of NLuc translocation was abrogated in an *icmSW* background. Furthermore, analysis using a bacterial two hybrid system supported direct interaction of the NLuc with IcmSW T4SS chaperones. In particular, the interaction of the N-terminus of NLuc with IcmW and also the inner membrane T4SS component, DotF, the latter previously used as bait to identify the Sid family of translocated effectors (44), was suggested by loss of interaction in N-terminal NLuc deletions.

Based on the crystal structure of NLuc, previously solved to 1.95 Å (45), the N-terminus is predicted to be surface exposed, potentially allowing it to interact directly with T4SS translocation machinery. In particular, we noted enrichment in negatively charged amino acids in the N-terminus, reminiscent of previously identified C-terminal translocation signals. However, specific replacement of these negatively charged amino acids either singly or in combination with uncharged amino acids or random scanning mutagenesis of each single N-terminal amino acid position, had only partial or no effect on translocation, suggesting a potential secondary role of the primary amino acid sequence in the N-terminal secretion signal. In contrast, disruption of the existing α-helical secondary structure, as predicted by RaptorX, either through amino acid deletion or multiple amino substitutions, almost completely abolished translocation, with rare exception. AlphaFold2 also predicted abolition of α-helical secondary structure in amino acid deletion mutants associated with translocation defects; in contrast, it predicted preservation of α-helical structure in variants with multiple amino acid substitutions including those defective in translocation. However, the ability of AlphaFold2 to accurately predict structural alterations associated with missense mutations has been questioned based on weighting of its algorithm towards recapitulation of existing PDB structures (46). Therefore, taking the mutagenesis data and structural predictions together, we suggest that surface-exposed α-helical secondary structure is critical to recognition by IcmSW-associated T4SS machinery.

We initially considered use of NLuc in one-step assays with a simple addition of luciferase substrate in the presence of saponin detergent, without need for extraction of eukaryotic cytoplasm and physical separation from infecting organisms. However, *Legionella* proved relatively permeable to commonly used furimazine substrate, perhaps related to *Legionella’s* overall increased permeability to small molecules (e.g., gram-positive antibiotics) compared with *Enterobacteriales* (47). Fortunately, less permeable coelenterazine analogs were identified (Table S1) that proved compatible with use in one-step T4SS translocation assays in a high throughput screening format. Furthermore, NLuc provided again an excellent fusion partner with GFP11, supporting its use as a translocation signal in an alternative, one single-step translocation assay based on detection of sfGFP in host cells. Although sfGFP assays have been previously used in translocation assays (48–50), a particularly exciting aspect of our bi-functional NLuc-based sfGFP constructs for high throughput screening drug discovery is that orthogonal confirmation of sfGFP results can be performed in the same screen wells using the alternative nanoluciferase readout.

Taken together, our data support use of NLuc-based T4SSb translocation assays. NLuc itself also provides a system for further exploration of the nature of T4SS secretion signals which likely include critical secondary structural motifs.

## Methods & Materials

### Bacterial strains and plasmids

Bacterial strains, plasmids, and eukaryotic cell lines used in this study are listed in Table 1. *Escherichia coli* NEB-5α (New England Biolabs, Beverley, MA) was grown in Luria broth (LB) medium (BD, NJ). *Legionella pneumophila* Lp02 *flaA* and Lp03 *flaA* strains were grown on buffered charcoal yeast extract (BCYE) medium supplemented with 100μg/mL of thymidine (51).

### Cells and materials

Murine J774A.1 (ATCC TIB-67, American Type Culture Collection, Manassas, VA) macrophage cell lines were cultivated in RPMI 1640 (Thermo Fisher Scientific, MA) with 9% heat-inactivated, iron-supplemented calf serum (GemCell, Gemini Bio-Products, CA) in a humidified 5% CO_2_ atmosphere at 37 °C. Furimazine was from Promega, catalogue N-110 (Madison, WI) or from AOBIOUS (Gloucester, MA) for bacterial permeability studies; coelenterazine, coelenterazine f, h, cp, fcp, hcp, i, ip, and n were from Biotium (Fremont, CA); and coelenterazine e, e-f, v, and 400a were from Nanolight Technology (Pinetop, AZ). Coelenterazine analogs were dissolved at 250 μg/ mL in ethanol. Prior to use, 1 μL of each substrate solution was added to 1 mL of PBS or RIPA lysis buffer (Cell Signaling Technology, Danvers, MA) for NLuc experiments. Fumarizine (Promega) was used at a 1:1000 dilution from purchased stock

### Generation of bacterial strains expressing NLuc fusion proteins

PCR products for gene or gene fragments were digested with appropriate restriction enzymes and cloned into similarly digested pXDC61 (EcoRI/KpnI) or pRetroX (BamHI/EcoRI). The NLuc sequence was amplified from *E. coli* strains harboring pNL1.1 (Promega, Madison, WI) (see primers in Table S4) and cloned into the vector, pXDC61 (52), with or without incorporation of a 3X FLAG tag immediately after the start codon or before the stop codon. For the construction of NLuc:RalF fusion proteins, the whole gene or the terminal 20 amino acids of RalF (amino acids 355-374) sequence was amplified from *L. pneumophila* and cloned into pXDC61-NLuc using restriction enzymes, KpnI and BamHI. Fusions of partial NLuc and TEV or TetR were constructed from corresponding gBlocks (Integrated DNA Technologies, Coralville, IA) and cloned into pXDC61. GFP11 fused with partial RalF or NLuc was cloned into pXDC61 using its EcoRI and KpnI restriction sites. GFPopt (synthesized as a gBlock, Integrated DNA Technologies) was amplified with indicated primers and cloned into the pRetroX vector using restriction enzymes, BamHI/EcoRI to create pRetroX-GFPopt. Plasmids were transformed into *E. coli* NEB-5α and/or *L. pneumophila* by electroporation as previously described (52), or from GFPopt transfected into J774A.1 cells as described below.

### NLuc Mutagenesis

A single-site saturation mutant library of NLuc was prepared using degenerate PCR primers in which the degenerate codon NNK was substituted for each amino acid position from V2 through D11 of NLuc (N indicates any nucleotide; K represents G or T). In addition, a multi-site saturation mutagenesis library was prepared in which all codons from V2 to W12 were replaced by NNK. PCR products were digested with EcoRI and KpnI enzymes, ligated into the similarly digest pXDC61, and then transformed into chemically competent *E. coli* NEB-5α. Representative plasmids clones were sequenced to identify specific amino acid substitutions, and select clones were transformed into *L. pneumophila* for translocation assays.

### Bacterial Two Hybridization Assay

The Bacterial Two Hybrid system (BACTH) was used to investigate protein-protein interaction between NLuc and Legionella Icw/Dot proteins. The readout of BATCH relies on the interaction of one partner protein fused with bacteriophage λ CI interacting with a second protein fused with the α subunit of *E. coli* RNA polymerase as previously described (27). Co-localization of λ CI and α RNP through binding together of the protein partners activates transcription of a *lacZ* reporter containing an upstream λCI-binding site. This system includes two plasmids, pACλCI, encoding λCI protein and pBRα, encoding the RNA polymerase α subunit protein under the control of the lac *UV5* promoter. Using primers designed to insert amplified proteins into the NotI/BamHI restriction sites of each plasmid, the *icmW*, *icmS*, and *dotF* open reading frames were cloned into the MCS of pACλCI. NLuc; NLuc16, missing the fifteen N-terminal amino acids of NLuc; RalF; and SidD genes were cloned into the multicloning site of pBRα plasmid. Indicated plasmid pairs were introduced into reporter strain FW102-OL2-62 for readout of lacZ reporter expression quantified in Miller units (53) after induction of protein expression with the indicated IPTG concentration. The fusion of λCI with the physiologically non-relevant β-flap subunit of *E. coli* RNA polymerase was used as a negative control.

### Macrophage Infection and Translocation Assays

One day before infection, the murine macrophage cell line, J774A.1 (American Type Culture Collectin, Manassas, VA) or macrophages transfected with GFP (as described below), cultured in RPMI medium (Fisher Scientific, Waltham, MA) supplemented with 9% iron-supplemented calf serum (Gemini Bio-Products, West Sacramento, CA), was seeded into white 96-well or clear 6-well tissue culture plates in the same medium. *L. pneumophila* grown on BCYE plate with selection for expression vector (5 μg/mL of chloramphenicol) and thymidine (100 μg/mL) was harvested by resuspending in saline, centrifugation at 9,300 x g for 2 min to pellet bacteria, followed by washing twice with PBS. The bacterial cells were re-suspended in RPMI supplemented with 100 µg/mL thymidine and used to infect J774A.1 cells at a multiplicity of infection (MOI) of 1 for 24 h or 10 for 6 h at 37°C in 5% CO_2_, as specified, in the presence of 0.5 mM of IPTG to induce expression of constructs.

At indicated time points, infected J774A.1 cells were washed 2x with PBS and lysed in PBS containing 0.2 % saponin on ice for 1 hr. The lysates were then centrifuged at 13,400 x g for 15 min. Supernatants, containing eukaryotic cytoplasm, were assayed for translocated NLuc activity using furimazine (Promega, Madison, WI) according to the manufacturer’s instructions or by western blot as described below. For one-step NLuc assays, infected J774A.1 cells in 100 µL well volume were lysed by addition of 50 µL PBS containing 0.6% saponin, and furimazine or indicated coelenterazine analogs. Luminescence was then read on a TECAN Infinite M1000 microplate reader (TECAN, Morrisville, NC). Green fluorescence, resulting from *in vivo* assembly of superfolder GFP, was measured on a TECAN Infinite M1000 plate reader using 485 nm excitation and 510 nm emission wavelengths.

### GFPopt-expressing J774A.1 Cells

Gryphon packaging cells (Allele Biotech, San Diego, CA) were transfected with 10 μg of pRetroX-GFPopt DNA using Lipofectamine LTX (ThermoFisher Scientific, Waltham, MA). After three days, supernatant was collected and used for transfection of J774A.1 cells. After a 3-day incubation, cells were replated in 6-well plates with RPMI medium containing 1mg/mL G-418 (Sigma-Aldrich, St. Louis, MO). As we described previously (52), individual colonies expressing GFP-opt were expanded in RPMI containing 500 μg/mL G-418 until use in experiments.

### Western blotting

After infection for indicated times, J774A.1 lysates were pelleted by centrifugation at 13,400 x g for 15 min at 4°C and denatured at 95 °C for 5 min in SDS-PAGE sample buffer containing 5% β-mercaptoethanol. Bacterial pellets or saponin extracts from equivalent cell numbers were separated by SDS-PAGE. After electrophoresis, proteins were transferred to a nitrocellulose membrane (Bio-Rad, Hercules, CA). Membranes were blocked with 5% non-fat dried milk in TBS plus 0.1% Tween 20 (TBST), incubated in primary and secondary antibodies diluted in 5% non-fat dried milk in TBST for 1 h each, and then washed with TBST. Primary antibodies (murine monoclonal ANTI-FLAG M2 antibody, Sigma-Aldrich #F1804; goat anti-GFP #600-101-215, Rockland Immunochemicals, Limerick, PA; and Mouse monoclonal anti-NanoLuc antibody, Promega #N7000) and secondary antibodies (goat anti-Mouse IgG (H+L) cross-adsorbed secondary antibody, HRP, ThermoFisher, #31432; and rabbit anti-Goat IgG (H+L) secondary antibody, HRP, ThermoFisher #31402) were used at a 1:1,000 or 1: 5,000 dilutions, respectively. Western blot signals were visualized by chemiluminescence using the SuperSignal™ West Femto Maximum Sensitivity Substrate (ThermoFisher Scientific) using a ChemiDoc (Bio-Rad, Hercules, CA).

### Confocal Microscopy

To observe *in vivo* assembly of sfGFP in J774A.1, GFPopt-expressing J774A.1 cells were plated in 12-well plates containing 1.5-thickness 12 mm round coverslips (Warner Instruments, Hamden, CT) at a density of 1 x 10^5^ cells per well. After 24 h, J774A.1 cells were infected with *L. pneumophila* expressing GFP11, GFP11::pRalF, or NLuc::GFP11 at a MOI of 1. Plates were immediately centrifuged at 930 x g for 10 min and then incubated at 37 °C with 5 % CO_2_. At 24 h post-infection, coverslips were fixed with 4 % formaldehyde in PBS for 30 minutes at room temperature, permeabilized with 0.3 % Tritox X-100 in PBS for 15 minutes, mounted in ProLong Gold with DAPI (Invitrogen, Carlsbad, CA), and visualized with a LSM 880 confocal microscope (Carl Zeiss Microscopy, White Plains, NY) using eGFP and DAPI settings for image collection. Image analysis was performed using Zen 2.1 software.

### Membrane permeability index determination

*L. pneumophila* Lp02 was grown overnight in AYET medium, washed twice with PBS, and resuspended in PBS at ∼1E8 bacteria/mL. 1.0 μL of each coelenterazine substate (resuspended in 250 μg/mL of EtOH) was added to 1 mL of PBS or RIPA cell lysis buffer. 50 μL of substrate solution and 50 µL of bacteria were combined in 96-weell black microtiter plates for luciferase determination on a TECAN Infinite M1000 microplate reader. Bacterial cell permeability was calculated based on the ratio of luminescence for bacteria resuspended in PBS compared to bacteria suspended in RIPA buffer. Data were normalized to results obtained for purified nanoluciferase (Nanolight Technologies, Pinetop, AZ), 20 ng per assay well, to account for effects of RIPA buffer on luciferase activity.

## Acknowledgments

This work was supported by the National Institute of Allergy and Infectious Diseases of the National Institutes of Health under award number R01AI099122 to J.E.K. The content is solely the responsibility of the authors and does not necessarily represent the official views of the National Institutes of Health. We thank TECAN for use of the Infinite Pro M1000. TECAN had no role in study design, manuscript preparation, or decision to publish. We thank Joe Vogel (Washington University) for isogenic *icmW*, *icmS*, *icmSW*, and *lvgA* and complemented *L. pneumophila* strains. We thank Thea Brennan-Krohn and Jessica Ross for critical reading of the manuscript.

## Supplementary Figure Legends

**Figure S1.**
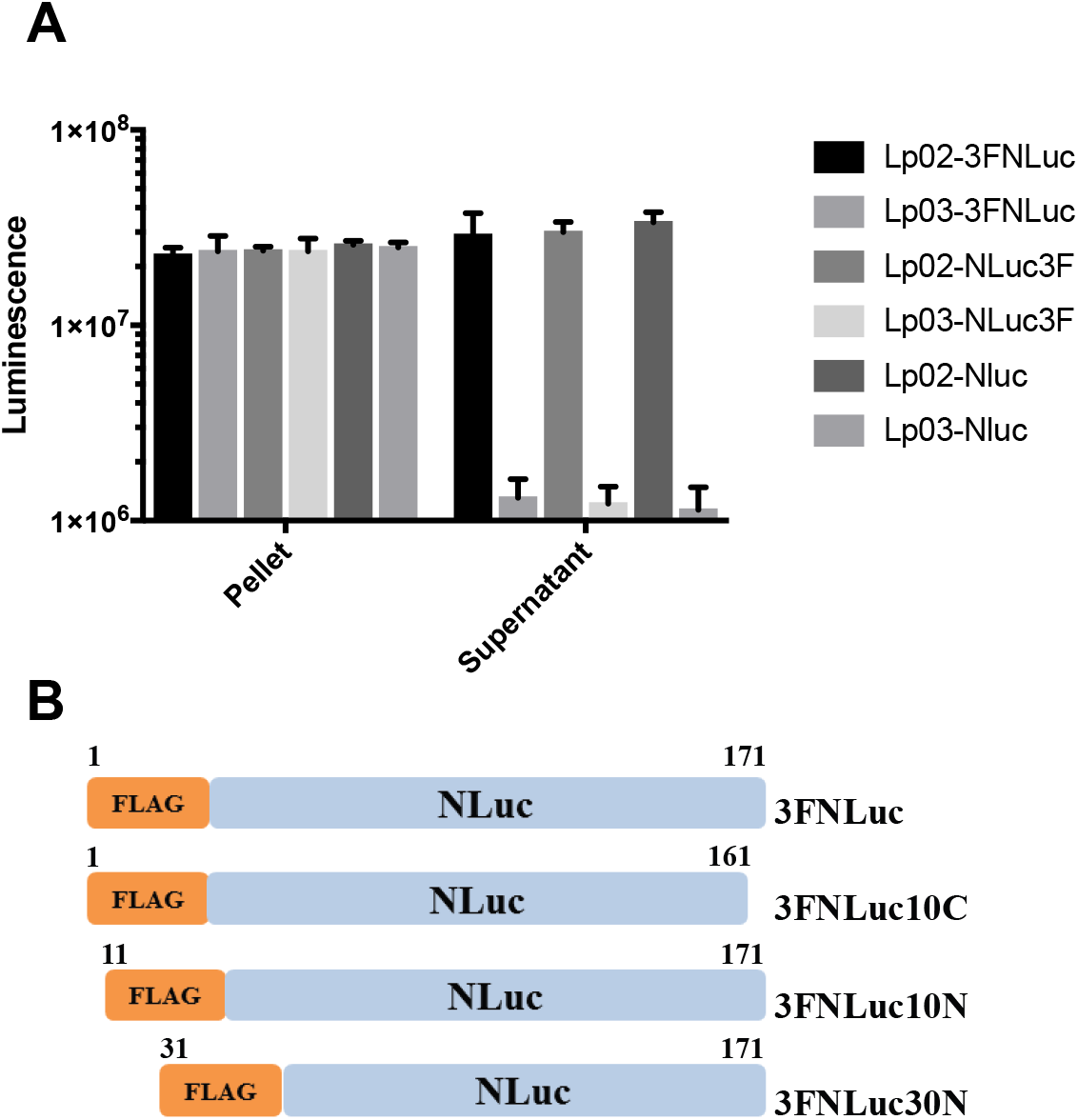
Identification of an N-terminal T4SS-dependent translocation signal in NLuc. **(A)** Luminescence of translocated NLuc extracted from eukaryotic cytoplasm (supernatant) and insoluble pellet 6 h post infection of J774A.1 cells infected with *L. pneumophila* Lp02 (wt) and Lp03 (*dotA*) expressing NLuc with an N-terminal (3FNluc) or C-terminal (NLuc3F) 3X-FLAG tags. Translocation of NLuc is evident and was not blocked by the 3X-FLAG in either an N-terminal or C-terminal position. Plotted are the mean and standard deviations of measurements from at least three biological replicates. **(B)** Illustration of NLuc deletion constructs used to establish location of an intrinsic translocation signal. Shown are wild type 3FNLuc; 3FNLuc10C (deletion of 10 C-terminal amino acids); 3FNluc10N (deletion of 10 N-terminal amino acids); and 3FNluc30N (deletion of 30 N-terminal terminal amino acids). As shown in Fig. 2B and 2C, T4SS-dependent translocation of constructs with both N-terminal deletions is abolished.

**Figure S2.**
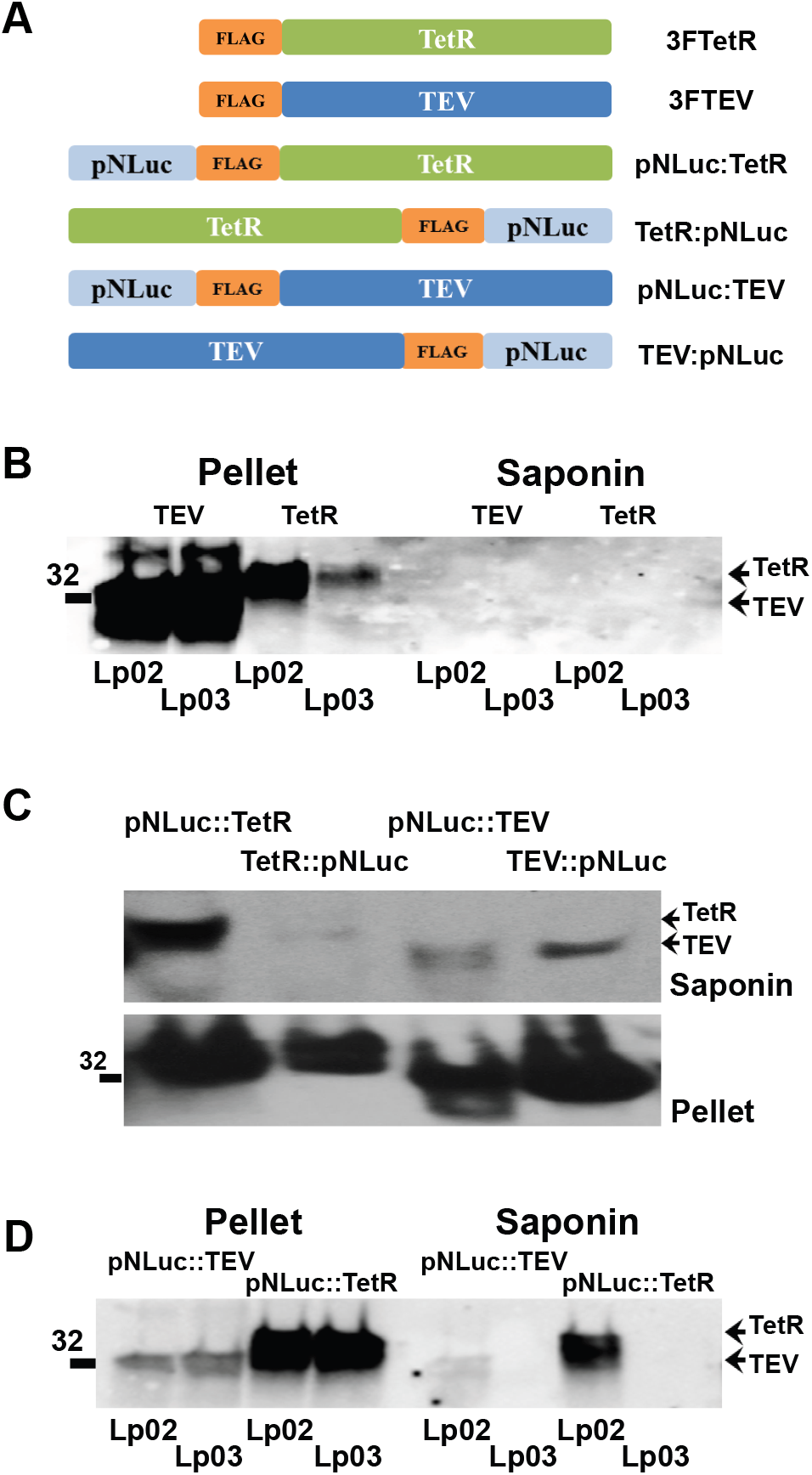
The N-terminus of NLuc is necessary and sufficient for T4SS-dependent translocation. **(A)** Structure of fusions of the 30 N-terminal amino acids of NLuc (pNluc) to the N- and C-terminus of TetR and TEV protease proteins, respectively, with incorporation of a 3X-FLAG epitope to enable detection by western blot. Fusions of 3X-FLAG to the N-terminus of TetR and TEV in the absence of NLuc were used as controls. **(B)** Western blots of J774A.1 cells infected with Lp02 or Lp03 at an MOI of 10 in strains expressing TEV and TetR proteins with 3X FLAG at the N-terminus in the absence of an NLuc translocation signal. At 6 h post infection, eukaryotic cytoplasm extract and insoluble pellet were separated by SDS-PAGE, and western blots probed with anti-FLAG tag antibody. TEV and TetR were only detected in the pellet fraction, indicating absence of an intrinsic translocation signal in these proteins. **(C)** Infections with strains expressing 3F-TEV or 3F-TetR fused to the first 30 amino acids of NLuc (pNluc) at the C- or N-terminus. All versions of TEV and TetR fused with pNLuc showed translocation into eukaryotic cytoplasm (supernatant), with the efficiencies of N-terminal and C-terminal fusions differing by protein partner. **(D)** Translocation only occurred when fusion proteins were expressed in T4SS-competent Lp02, but not in T4SS-incompetent, Lp03.

**Figure S3.**
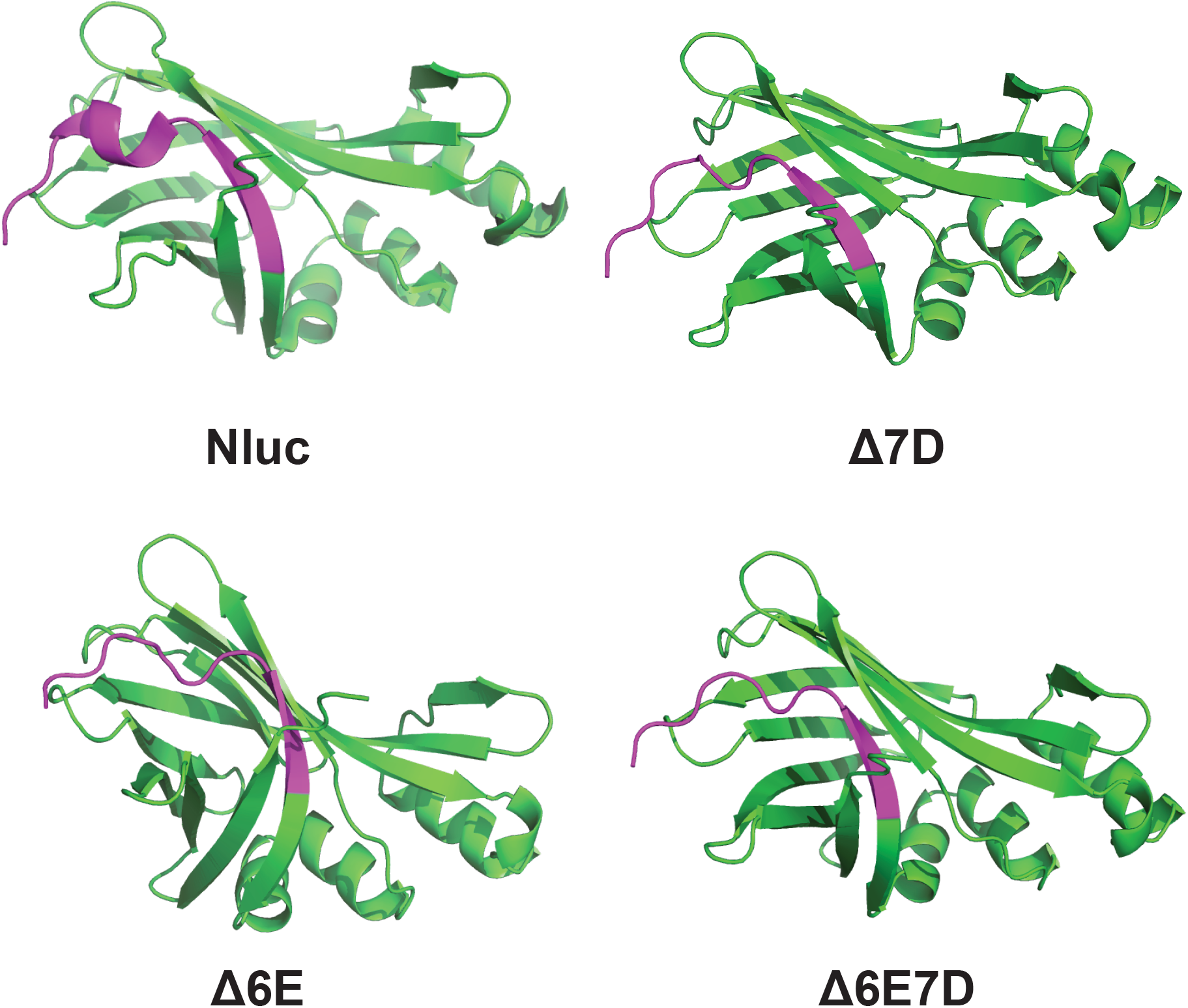
AlphaFold2 predictions of mutant NLuc structures. AlphaFold2 CoLab (36) predictions of the structure of NLuc, NLuc1Δ6E, NLuc1Δ7D, and NLuc1Δ6E1Δ7D. The surface-exposed, N-termini (M1-W12) are colored magenta. In the three deletion mutants, the N-terminal α-helix, present in the structure predicted by AlphaFold2 and the solved structure of NLuc (PDB 5IBO), is now an unstructured region.

**Figure S4.**
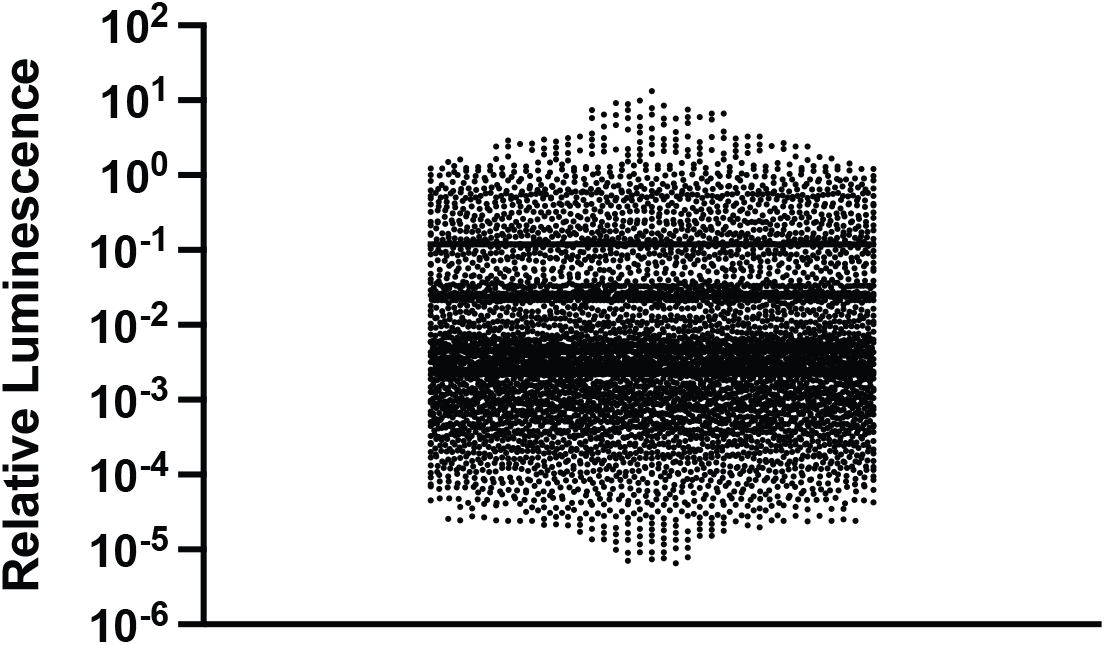
Luminescence associated with random mutagenesis of the N-terminus of NLuc. Approximately seven thousand random mutants of the NLuc N-terminus, inclusive of amino acids V2 to W12, were generated using a degenerate, primer randomization strategy and were tested for preservation of luminescence when expressed in *E. coli K-12*. Shown are the luminescence of analyzed clones normalized to wild type NLuc. Approximately 30 clones with preserved and/or elevated luminescence were further tested in *L. pneumophila* for T4SS-dependent translocation, as described in the main body of the manuscript.

**Figure S5.**
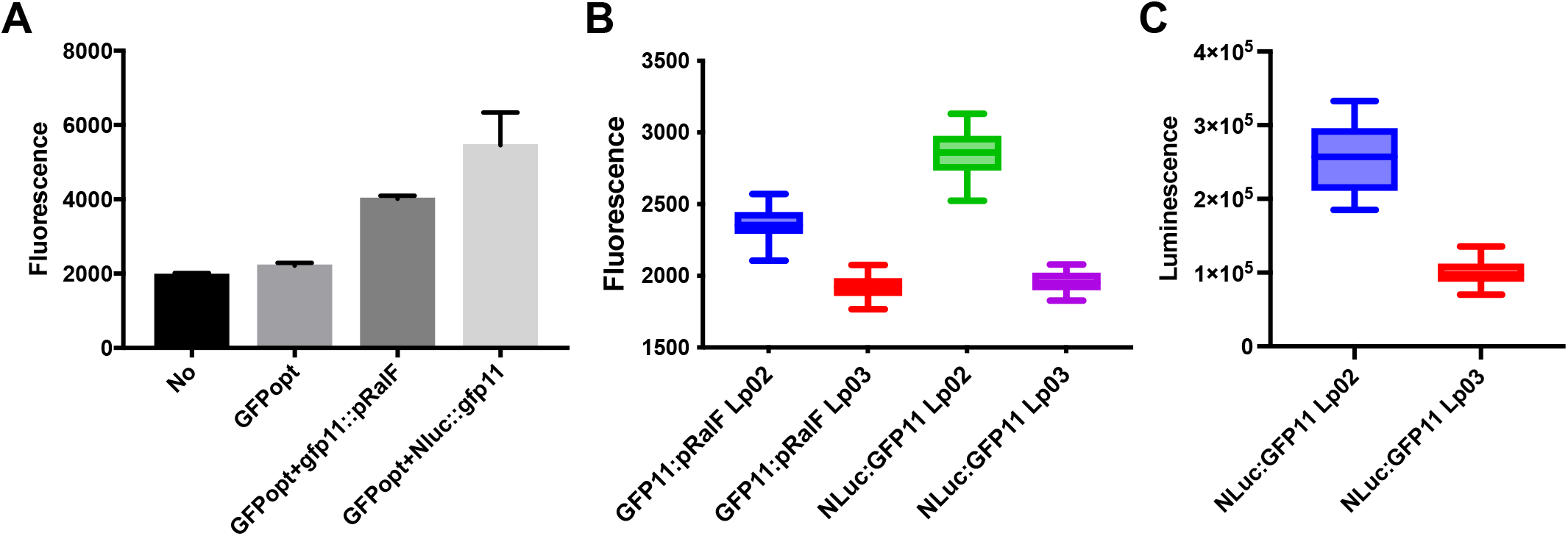
Development of a split GFP high throughput T4SS assay using an NLuc translocation signal. **(A)** The non-fluorescent GFP11 fragment and GFPopt protein can combine to yield fluorescent superfolder GFP (sfGFP). To determine whether this assembly remained functional when T4SS translocation signals were attached to the small GFP11 fragment, the fluorescent signal from combination of plasmids expressing GFP11 fused with NLuc (GFP11:NLuc) or partial RalF (GFP11:pRalF) and GFPopt were co-expressed in *E. coli* K-12. Bars represent the mean and standard deviation for three independent experiments performed in duplicate after overnight incubation and induction of protein expression with 0.5 mM IPTG. **(B)** The ability to use this system in a high throughput screening format to quantify T4SS-dependent translocation was assessed by infecting J774A.1 cells expressing GFPopt with *L. pneumophila* strains producing GFP11:pRalF or NLuc:GFP11. Fluorescent intensity was assayed in white 96-well plates, and mean and standard deviations determined from the combined data from two independent experiments, each performed with 40 technical replicates. Data were used to determined Z’ described in the results section. **(C)** NLuc:GFP11 also provided a single-step luciferase readout in combination with coelenterazine h. Mean and standard deviations were determined from the combined data from two independent experiments, each performed with ten technical replicates. Notably, through infecting J774A.1 GFPopt host cells with bacteria expressing NLuc:GFP11, the same NLuc:GFP11 construct can provide orthogonal sfGFP fluorescence- and nanoluciferase-based translocation readouts in the same high throughput screening wells, thereby presumably significantly reducing the burden of time-consuming secondary assays for screening hit confirmation.

**Table S1.** Effects of Single Amino Acid Substitutions in V2 through D11 of Nluc on Luciferase Activity and Translocation.

|  | Luminescence intensity<sup>a, b</sup> |  | Translocation efficiency<sup>a</sup> |  | Average translocation efficiency | Normalized translocation<sup>c</sup> |
| --- | --- | --- | --- | --- | --- | --- |
| NLuc wt |  |  |  |  |  |  |
| Lp02fla | 1.00 | 1.00 | 1.00 | 1.00 | 1.00 | 100% |
| Nluc wt |  |  |  |  |  |  |
| Lp03fla | 1.21 | 1.05 | 0.08 | 0.16 | 0.12 | 0% |
| V2Lys | 2.04 | 1.69 | 1.43 | 1.48 | 1.45 | 151% |
| V2Ala | 0.51 | 1.42 | 1.23 | 1.40 | 1.31 | 136% |
| V2Ser | 0.29 | 0.54 | 0.83 | 0.88 | 0.86 | 84% |
| V2Trp | 0.74 | 0.61 | 1.42 | 1.55 | 1.48 | 155% |
| V2Gly | 1.17 | 0.98 | 1.27 | 1.47 | 1.37 | 142% |
| V2Asn | 1.49 | 1.24 | 0.57 | 1.39 | 0.98 | 98% |
| V2His | 4.72 | 1.93 | 1.42 | 1.28 | 1.35 | 140% |
| V2Phe | 2.22 | 1.85 | 1.30 | 1.43 | 1.37 | 142% |
| V2Ile | 0.00 | 0.03 | 0.65 | 1.24 | 0.94 | 94% |
| V2Met | 1.11 | 0.93 | 1.23 | 1.36 | 1.30 | 134% |
| V2Tyr | 3.60 | 1.40 | 1.25 | 1.30 | 1.28 | 132% |
| V2Glu | 0.20 | 0.57 | 2.35 | 1.71 | 2.03 | 217% |
| V2Arg | 0.52 | 0.83 | 1.66 | 1.63 | 1.65 | 174% |
| V2Asp | 0.18 | 0.35 | 2.59 | 1.55 | 2.07 | 222% |
| V2Cys | 1.14 | 0.95 | 1.14 | 1.27 | 1.21 | 124% |
| V2Pro | 0.02 | 0.12 | 1.03 | 1.47 | 1.25 | 129% |
| V2Thr | 2.71 | 1.26 | 1.18 | 1.31 | 1.25 | 128% |
| V2Leu | 2.79 | 1.33 | 1.19 | 1.16 | 1.17 | 120% |
| V2Gln | 1.16 | 0.96 | 1.32 | 1.24 | 1.28 | 132% |
| F3Ala | 1.67 | 1.09 | 0.55 | 1.16 | 0.85 | 83% |
| F3Ile | 2.07 | 1.12 | 1.11 | 1.72 | 1.42 | 147% |
| F3Asn | 2.51 | 1.29 | 0.57 | 0.50 | 0.54 | 47% |
| F3Glu | 1.26 | 1.05 | 0.66 | 1.02 | 0.84 | 82% |
| F3Gln | 0.72 | 0.80 | 0.41 | 1.28 | 0.84 | 82% |
| F3Met | 1.45 | 1.20 | 1.26 | 1.06 | 1.16 | 118% |
| F3Thr | 1.21 | 0.81 | 1.12 | 1.32 | 1.22 | 125% |
| F3Pro | 2.76 | 1.29 | 0.84 | 1.19 | 1.01 | 102% |
| F3Leu | 2.34 | 1.34 | 0.84 | 1.45 | 1.14 | 116% |
| F3Ser | 2.00 | 1.27 | 0.45 | 1.06 | 0.75 | 72% |
| F3Tyr | 1.65 | 0.97 | 1.07 | 1.32 | 1.20 | 122% |
| F3His | 8.13 | 2.77 | 1.12 | 0.96 | 1.04 | 105% |
| F3Lys | 1.57 | 1.31 | 0.98 | 1.05 | 1.02 | 102% |
| F3Arg | 0.86 | 1.71 | 0.61 | 0.96 | 0.78 | 75% |
| F3Cys | 3.82 | 1.18 | 0.53 | 0.67 | 0.60 | 55% |
| F3Val | 2.28 | 1.30 | 0.35 | 0.50 | 0.43 | 35% |
| F3Gly | 1.78 | 1.48 | 0.44 | 0.59 | 0.52 | 45% |
| F3Trp | 3.02 | 1.32 | 0.37 | 0.52 | 0.45 | 37% |
| F3Asp | 5.10 | 2.25 | 0.44 | 0.76 | 0.60 | 55% |
| T4Gly | 0.74 | 1.62 | 1.05 | 1.19 | 1.12 | 114% |
| T4Pro | 1.56 | 1.30 | 0.86 | 1.46 | 1.16 | 118% |
| T4Val | 1.51 | 1.25 | 0.82 | 0.83 | 0.82 | 80% |
| T4Cys | 0.01 | 0.10 | 1.32 | 1.01 | 1.16 | 118% |
| T4Leu | 1.58 | 1.21 | 0.83 | 0.97 | 0.90 | 88% |
| T4Ser | 0.87 | 0.72 | 1.07 | 0.98 | 1.03 | 103% |
| T4Trp | 7.10 | 2.91 | 1.30 | 1.45 | 1.37 | 142% |
| T4Asn | 9.61 | 3.00 | 1.42 | 1.43 | 1.42 | 148% |
| T4Lys | 9.75 | 3.11 | 1.44 | 1.57 | 1.50 | 157% |
| T4Gln | 0.70 | 0.58 | 1.09 | 1.24 | 1.16 | 119% |
| T4Asp | 2.53 | 1.61 | 1.45 | 1.33 | 1.39 | 145% |
| T4Arg | 4.87 | 2.05 | 1.18 | 1.20 | 1.19 | 122% |
| T4Met | 5.77 | 2.80 | 1.78 | 1.60 | 1.69 | 179% |
| T4Tyr | 6.85 | 2.70 | 1.47 | 1.12 | 1.29 | 133% |
| T4His | 1.34 | 1.11 | 0.88 | 1.03 | 0.95 | 95% |
| T4Phe | 1.24 | 1.03 | 0.90 | 0.92 | 0.91 | 90% |
| T4Ile | 0.60 | 0.80 | 0.98 | 1.22 | 1.10 | 111% |
| T4Ala | 1.17 | 0.98 | 1.08 | 1.09 | 1.09 | 110% |
| T4Glu | 0.98 | 0.82 | 1.58 | 1.12 | 1.35 | 140% |
| L5Arg | 0.00 | 0.02 | 1.38 | 1.28 | 1.33 | 138% |
| L5Gly | 1.25 | 1.04 | 0.76 | 0.63 | 0.69 | 65% |
| L5Trp | 1.58 | 1.32 | 1.23 | 1.15 | 1.19 | 122% |
| L5Thr | 1.56 | 1.30 | 1.64 | 1.52 | 1.58 | 166% |
| L5Gln | 1.48 | 1.13 | 0.96 | 0.83 | 0.90 | 88% |
| L5Glu | 1.44 | 1.10 | 0.86 | 0.84 | 0.85 | 83% |
| L5Ala | 1.12 | 0.94 | 0.82 | 0.84 | 0.83 | 80% |
| L5Lys | 1.58 | 1.32 | 2.82 | 1.28 | 2.05 | 220% |
| L5Ser | 0.08 | 0.04 | 1.23 | 1.52 | 1.37 | 143% |
| L5Cys | 1.06 | 0.88 | 0.82 | 1.15 | 0.99 | 98% |
| L5Asp | 0.05 | 0.10 | 1.01 | 1.01 | 1.01 | 102% |
| L5Asn | 0.44 | 0.37 | 0.89 | 1.02 | 0.95 | 95% |
| L5Phe | 0.07 | 0.06 | 0.89 | 0.91 | 0.90 | 89% |
| L5Ile | 3.24 | 1.70 | 1.06 | 1.09 | 1.08 | 109% |
| L5Met | 1.74 | 1.25 | 1.21 | 1.11 | 1.16 | 118% |
| L5Val | 0.15 | 0.06 | 0.86 | 0.76 | 0.81 | 78% |
| L5His | 0.06 | 0.15 | 0.85 | 0.98 | 0.92 | 91% |
| L5Tyr | 0.09 | 0.07 | 0.99 | 1.24 | 1.11 | 113% |
| L5Pro | 0.35 | 0.19 | 0.83 | 0.88 | 0.86 | 84% |
| E6Cys | 3.24 | 1.69 | 1.37 | 0.94 | 1.16 | 118% |
| E6Val | 0.39 | 0.82 | 0.82 | 0.57 | 0.70 | 66% |
| E6Gly | 0.20 | 0.97 | 0.38 | 0.50 | 0.44 | 36% |
| E6Trp | 1.28 | 1.26 | 0.93 | 0.76 | 0.85 | 82% |
| E6Arg | 1.53 | 1.27 | 1.02 | 0.66 | 0.84 | 82% |
| E6Pro | 1.72 | 1.13 | 0.87 | 0.51 | 0.69 | 64% |
| E6Gln | 0.59 | 1.49 | 1.01 | 0.65 | 0.83 | 81% |
| E6Leu | 0.75 | 1.62 | 1.16 | 0.81 | 0.99 | 99% |
| E6Ser | 2.68 | 1.23 | 1.07 | 0.73 | 0.90 | 89% |
| E6Ala | 0.13 | 0.81 | 0.69 | 0.36 | 0.52 | 46% |
| E6Ile | 0.86 | 0.92 | 0.96 | 0.63 | 0.80 | 77% |
| E6His | 0.76 | 1.64 | 0.79 | 0.46 | 0.63 | 57% |
| E6Met | 0.89 | 0.94 | 1.33 | 0.99 | 1.16 | 118% |
| E6Lys | 4.70 | 0.91 | 1.03 | 0.68 | 0.86 | 84% |
| E6Tyr | 3.84 | 1.19 | 1.36 | 1.01 | 1.19 | 121% |
| E6Thr | 0.79 | 1.66 | 1.34 | 0.98 | 1.16 | 118% |
| E6Asn | 3.59 | 0.99 | 1.47 | 1.10 | 1.29 | 133% |
| E6Asp | 2.99 | 1.49 | 1.00 | 0.63 | 0.82 | 79% |
| E6Phe | 2.57 | 1.14 | 1.14 | 0.77 | 0.95 | 95% |
| D7Thr | 3.40 | 1.83 | 0.67 | 0.54 | 0.61 | 55% |
| D7Trp | 1.75 | 1.06 | 0.54 | 0.54 | 0.54 | 47% |
| D7Arg | 0.45 | 0.78 | 0.38 | 0.41 | 0.40 | 31% |
| D7Cys | 2.35 | 1.05 | 0.85 | 0.68 | 0.77 | 73% |
| D7Ala | 3.12 | 1.90 | 0.54 | 0.72 | 0.63 | 58% |
| D7Leu | 0.54 | 0.75 | 0.46 | 0.41 | 0.44 | 36% |
| D7Asn | 2.28 | 1.39 | 1.52 | 1.40 | 1.46 | 153% |
| D7Gly | 1.17 | 0.98 | 0.55 | 0.50 | 0.52 | 46% |
| D7Val | 1.16 | 0.97 | 0.34 | 0.42 | 0.38 | 29% |
| D7Pro | 2.41 | 1.00 | 0.63 | 0.50 | 0.57 | 51% |
| D7Phe | 0.00 | 0.01 | 0.41 | 0.35 | 0.38 | 29% |
| D7Ile | 0.04 | 0.09 | 0.77 | 0.65 | 0.71 | 67% |
| D7Met | 0.18 | 0.05 | 0.48 | 0.36 | 0.42 | 34% |
| D7Ser | 0.00 | 0.03 | 0.42 | 0.49 | 0.46 | 38% |
| D7Tyr | 0.00 | 0.01 | 0.46 | 0.54 | 0.50 | 43% |
| D7His | 0.32 | 0.16 | 0.46 | 0.34 | 0.40 | 31% |
| D7Gln | 1.55 | 0.29 | 0.53 | 0.41 | 0.47 | 40% |
| D7Lys | 1.23 | 1.02 | 0.81 | 0.79 | 0.80 | 77% |
| D7Glu | 0.00 | 0.01 | 0.50 | 0.37 | 0.44 | 36% |
| F8Arg | 0.00 | 0.01 | 1.89 | 1.55 | 1.72 | 182% |
| F8Ser | 0.02 | 0.01 | 0.86 | 0.66 | 0.76 | 73% |
| F8Pro | 0.01 | 0.01 | 1.34 | 1.37 | 1.36 | 141% |
| F8Gly | 0.02 | 0.03 | 0.88 | 1.10 | 0.99 | 99% |
| F8Gln | 0.01 | 0.04 | 0.99 | 0.76 | 0.87 | 86% |
| F8Ala | 0.26 | 0.12 | 0.91 | 0.58 | 0.74 | 71% |
| F8Thr | 0.28 | 0.13 | 0.90 | 0.92 | 0.91 | 90% |
| F8Leu | 0.21 | 0.18 | 1.11 | 0.77 | 0.94 | 93% |
| F8Ile | 0.21 | 0.19 | 0.93 | 0.96 | 0.94 | 93% |
| F8Met | 0.18 | 0.15 | 1.19 | 1.26 | 1.22 | 126% |
| F8Tyr | 1.96 | 1.16 | 1.04 | 0.81 | 0.92 | 91% |
| F8His | 0.00 | 0.01 | 0.76 | 0.78 | 0.77 | 74% |
| F8Asn | 0.00 | 0.02 | 0.83 | 0.60 | 0.72 | 68% |
| F8Lys | 0.00 | 0.01 | 1.13 | 0.90 | 1.01 | 102% |
| F8Asp | 0.00 | 0.02 | 0.93 | 0.60 | 0.76 | 73% |
| F8Glu | 0.00 | 0.02 | 1.13 | 1.16 | 1.14 | 116% |
| F8Cys | 0.04 | 0.01 | 1.02 | 1.23 | 1.13 | 114% |
| F8Trp | 0.34 | 0.18 | 1.04 | 0.70 | 0.87 | 85% |
| F8Val | 0.63 | 0.53 | 0.98 | 1.01 | 1.00 | 100% |
| V9Ser | 3.00 | 1.49 | 1.44 | 1.31 | 1.37 | 143% |
| V9Asp | 2.78 | 1.31 | 1.23 | 1.10 | 1.17 | 119% |
| V9Glu | 1.34 | 1.12 | 0.99 | 1.04 | 1.02 | 102% |
| V9Gly | 0.00 | 0.08 | 2.16 | 1.80 | 1.98 | 211% |
| V9Phe | 0.80 | 0.27 | 0.96 | 0.60 | 0.78 | 75% |
| V9Thr | 5.25 | 2.37 | 1.46 | 1.10 | 1.28 | 132% |
| V9Cys | 1.31 | 1.69 | 1.78 | 1.36 | 1.57 | 165% |
| V9Pro | 0.64 | 0.73 | 1.54 | 1.18 | 1.36 | 141% |
| V9Arg | 1.46 | 1.02 | 1.72 | 1.36 | 1.54 | 162% |
| V9Leu | 0.97 | 0.50 | 1.00 | 0.86 | 0.93 | 92% |
| V9Ile | 2.16 | 1.30 | 0.97 | 1.02 | 0.99 | 99% |
| V9Met | 1.44 | 1.20 | 0.82 | 0.90 | 0.86 | 83% |
| V9Ala | 1.61 | 1.04 | 0.88 | 0.52 | 0.70 | 66% |
| V9Tyr | 0.00 | 0.01 | 0.64 | 0.72 | 0.68 | 63% |
| V9His | 0.52 | 0.13 | 0.76 | 0.82 | 0.79 | 76% |
| V9Gln | 3.85 | 2.21 | 0.87 | 0.73 | 0.80 | 77% |
| V9Asn | 0.66 | 0.55 | 1.31 | 0.95 | 1.13 | 115% |
| V9Lys | 0.04 | 0.04 | 0.91 | 0.99 | 0.95 | 95% |
| V9Trp | 0.44 | 0.17 | 1.01 | 0.65 | 0.83 | 81% |
| G10Ala | 4.32 | 1.60 | 1.08 | 0.95 | 1.01 | 102% |
| G10Arg | 0.15 | 0.12 | 0.89 | 0.56 | 0.73 | 69% |
| G10Cys | 1.45 | 1.20 | 1.00 | 1.05 | 1.03 | 103% |
| G10Phe | 4.03 | 1.36 | 1.24 | 1.11 | 1.18 | 120% |
| G10Leu | 0.43 | 0.35 | 0.99 | 1.23 | 1.11 | 112% |
| G10Trp | 0.25 | 0.11 | 0.86 | 0.53 | 0.69 | 65% |
| G10Glu | 0.55 | 0.35 | 1.24 | 1.48 | 1.36 | 141% |
| G10Val | 0.17 | 0.19 | 1.04 | 1.09 | 1.06 | 107% |
| G10Ile | 0.01 | 0.03 | 0.93 | 0.79 | 0.86 | 84% |
| G10Met | 0.02 | 0.03 | 0.90 | 1.14 | 1.02 | 102% |
| G10Ser | 0.38 | 0.12 | 0.86 | 0.92 | 0.89 | 87% |
| G10Pro | 0.00 | 0.03 | 0.79 | 1.19 | 0.99 | 98% |
| G10Thr | 0.00 | 0.01 | 0.96 | 0.81 | 0.88 | 87% |
| G10Tyr | 0.00 | 0.02 | 0.80 | 0.94 | 0.87 | 85% |
| G10His | 0.01 | 0.02 | 0.71 | 0.93 | 0.82 | 79% |
| G10Gln | 0.02 | 0.03 | 0.89 | 0.85 | 0.87 | 85% |
| G10Asn | 0.11 | 0.04 | 0.88 | 1.19 | 1.03 | 104% |
| G10Lys | 0.00 | 0.02 | 0.76 | 0.93 | 0.85 | 82% |
| G10Asp | 0.58 | 0.48 | 1.13 | 0.81 | 0.97 | 97% |
| D11Glu | 5.07 | 3.22 | 1.44 | 1.22 | 1.33 | 138% |
| D11Lys | 6.68 | 3.56 | 0.54 | 0.73 | 0.63 | 58% |
| D11Ala | 5.17 | 2.30 | 0.76 | 0.58 | 0.67 | 62% |
| D11Met | 0.69 | 0.57 | 0.95 | 1.29 | 1.12 | 114% |
| D11Thr | 3.31 | 2.75 | 0.91 | 1.11 | 1.01 | 101% |
| D11Arg | 4.75 | 2.95 | 0.62 | 0.82 | 0.72 | 68% |
| D11Cys | 6.62 | 3.51 | 1.06 | 1.18 | 1.12 | 114% |
| D11Gly | 4.35 | 3.62 | 1.07 | 1.14 | 1.11 | 112% |
| D11Leu | 4.48 | 2.73 | 1.20 | 0.88 | 1.04 | 104% |
| D11Phe | 0.00 | 0.00 | 1.08 | 1.01 | 1.05 | 105% |
| D11Ile | 0.01 | 0.00 | 0.95 | 0.81 | 0.88 | 86% |
| D11Ser | 0.01 | 0.01 | 1.07 | 1.33 | 1.20 | 123% |
| D11Pro | 0.00 | 0.02 | 1.00 | 0.88 | 0.94 | 93% |
| D11Val | 0.00 | 0.00 | 0.92 | 1.27 | 1.10 | 111% |
| D11Tyr | 0.00 | 0.00 | 1.08 | 1.26 | 1.17 | 120% |
| D11His | 0.00 | 0.03 | 1.04 | 0.85 | 0.95 | 94% |
| D11Gln | 0.07 | 0.10 | 0.87 | 1.20 | 1.03 | 104% |
| D11Asn | 0.67 | 0.56 | 1.00 | 1.28 | 1.14 | 116% |
| D11Trp | 0.13 | 0.21 | 0.98 | 0.85 | 0.92 | 91% |

<sup>a</sup>replicates are from two independent experiments (biological replicates), each infection with the indicated NLuc mutant performed in a well of a 6-well tissue culture plate.
<sup>b</sup>Luminescence readings are from bacterial pellets after saponin extraction of eukaryotic cytoplasm.
<sup>c</sup>normalized to translocation of wild type Nluc observed in Lp02 and Lp03 set at 100% and 0% respectively.

**Table S2.**
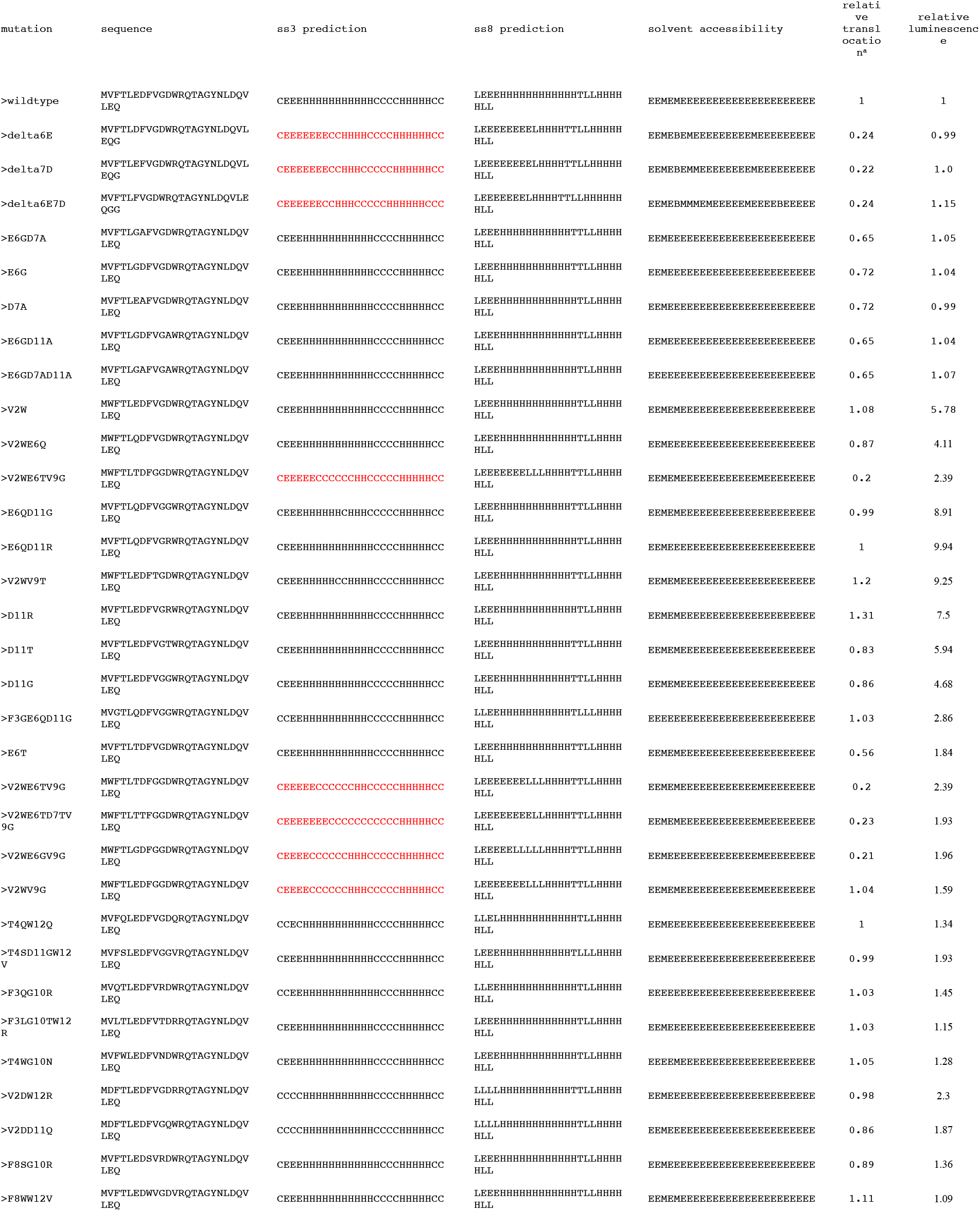
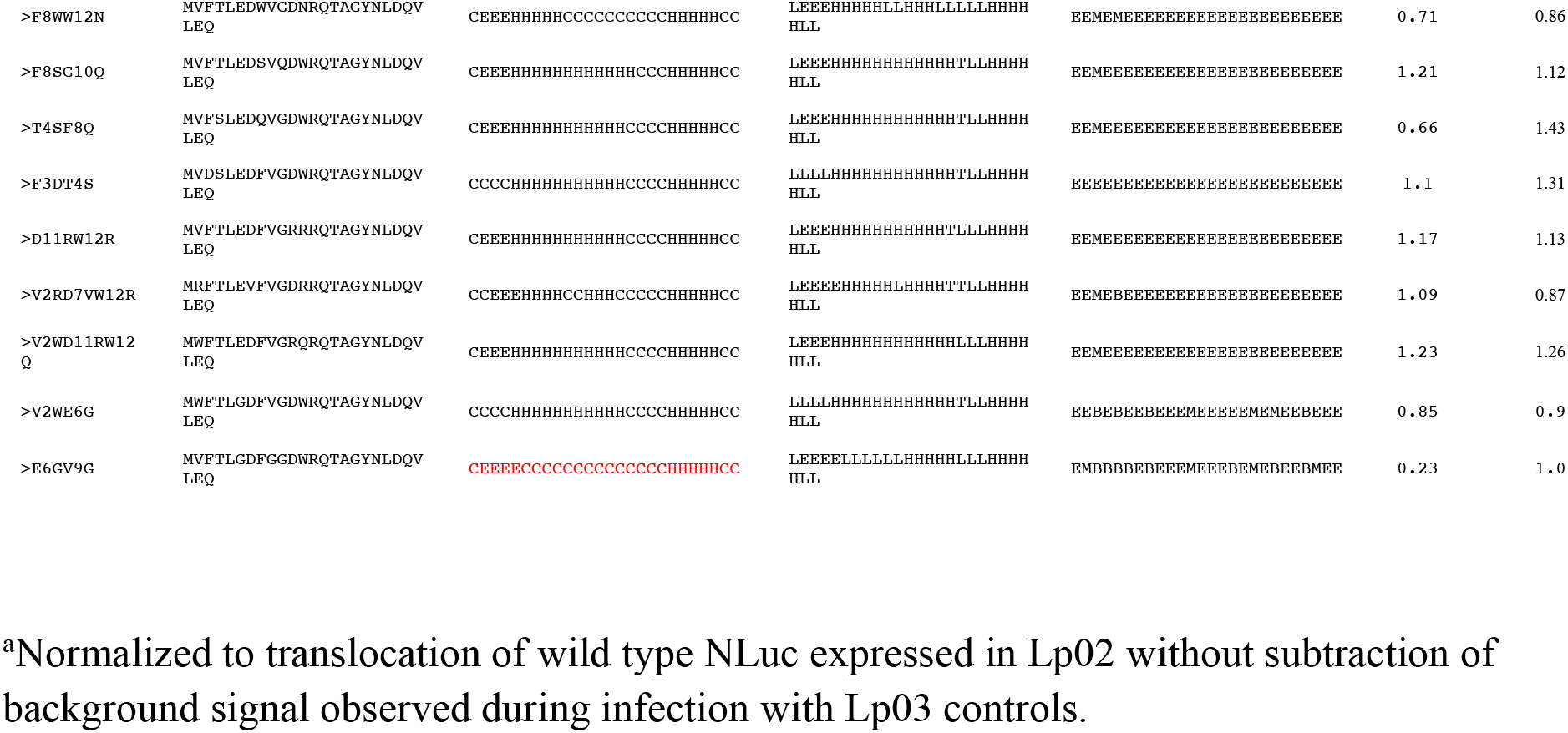
RaptorX Predicted Secondary Structure of the N-termini of Wild Type and Mutated Versions of Nanoluciferase.

**Table S3.** Relative permeability and luminesce of commercially available furimazine analogs.

| <u>analog</u> | <u>relative permeability<sup>a</sup></u> | <u>relative luminescence<sup>b</sup></u> |
| --- | --- | --- |
| furimazine | 3.1±0.3 | 1 |
| coelenterazine | 1.9±0.4 | 0.12 |
| 400a | 0.6±0.1 | 7.7 |
| cp | 0.4±0.1 | 0.09 |
| e | 1.3±0.1 | 0.62 |
| e-F | 3.3±0.4 | 0.69 |
| f | 4.9±0.3 | 0.21 |
| fcp | 1.0±0.1 | 0.53 |
| h | 0.4±0.1 | 2.8 |
| hcp | 5.3±0.7 | 0.12 |
| i | 0.7±0.0 | 1.6 |
| ip | 0.2±0.1 | 0.02 |
| n | 0.5±0.0 | 2.5 |
| v | 1.0±0.1 | 0.01 |

<sup>a</sup>relative permeability index was calculated as the ratio of luminescence from bacteria suspended in PBS to the luminescence of bacteria permeabilized in radioimmunoprecipitation (RIPA) buffer containing 1% NP-40 and 1% sodium deoxycholate. Results were normalized to activity of purified nanoluciferase added to PBS and RIPA alone, as RIPA buffer variably inhibited or stimulated activity of different nanoluciferase substrates up to 10-fold. The relative permeability ratios are the mean of values obtained from two independent experiments each with two technical replicates.
<sup>b</sup>luminescence signal relative to furimazine substrate based on testing an equivalent amount of Lp02 bacteria expressing nanoluciferase permeabilized in RIPA buffer, i.e., the denominator in the relative permeability index.

**Table S4.**
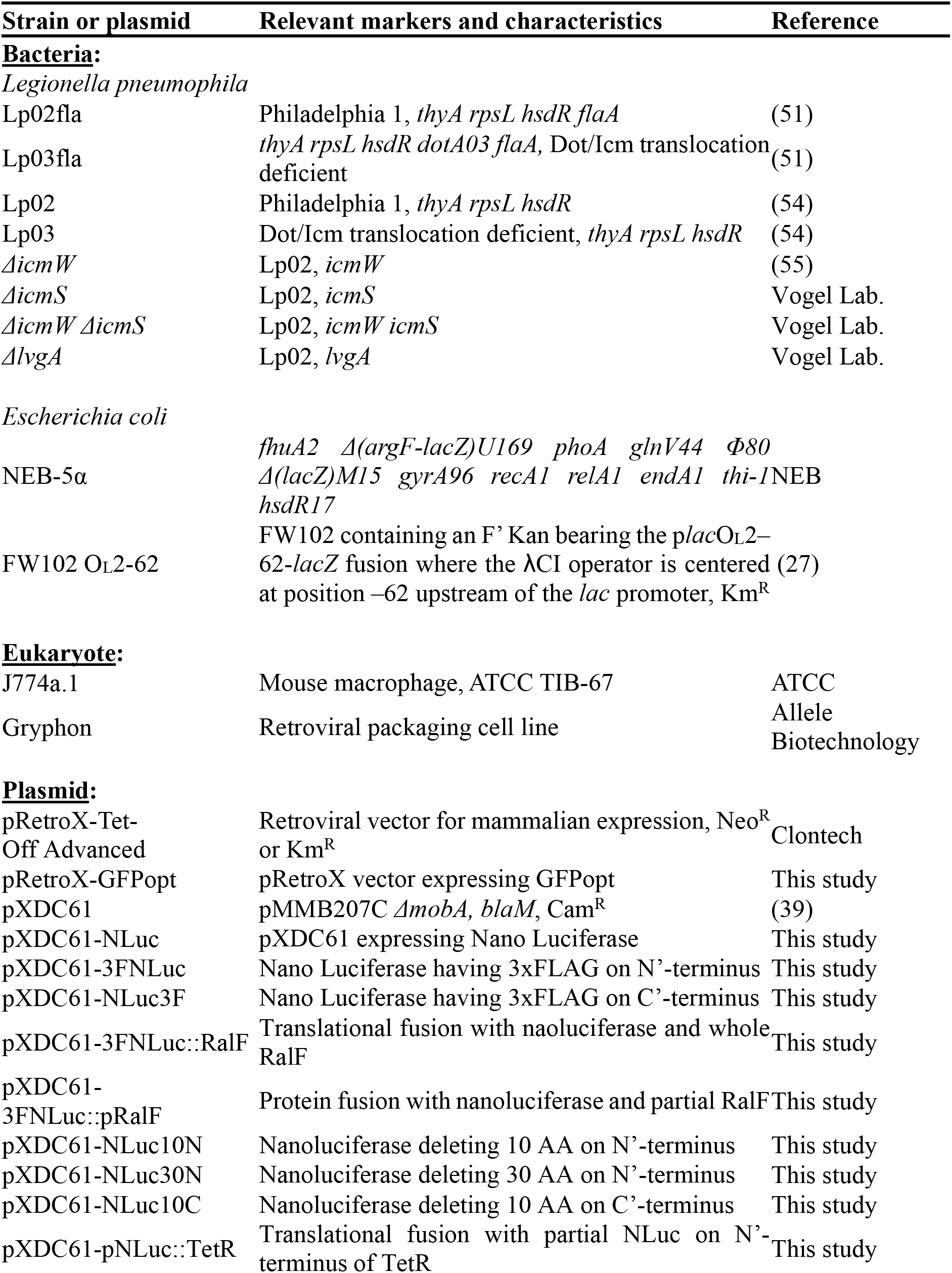
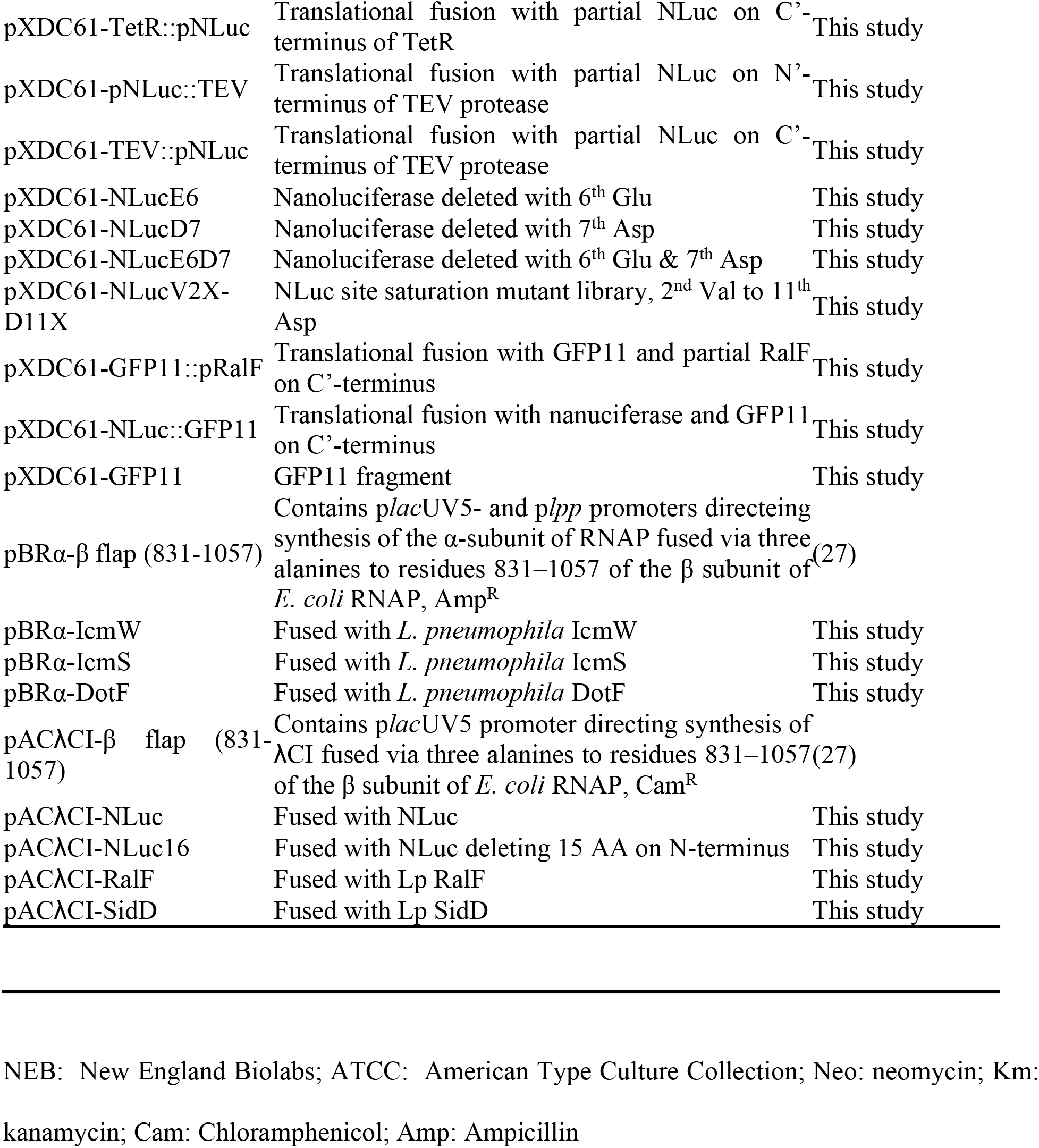
Bacterial strains and plasmids.

**Table S5.**
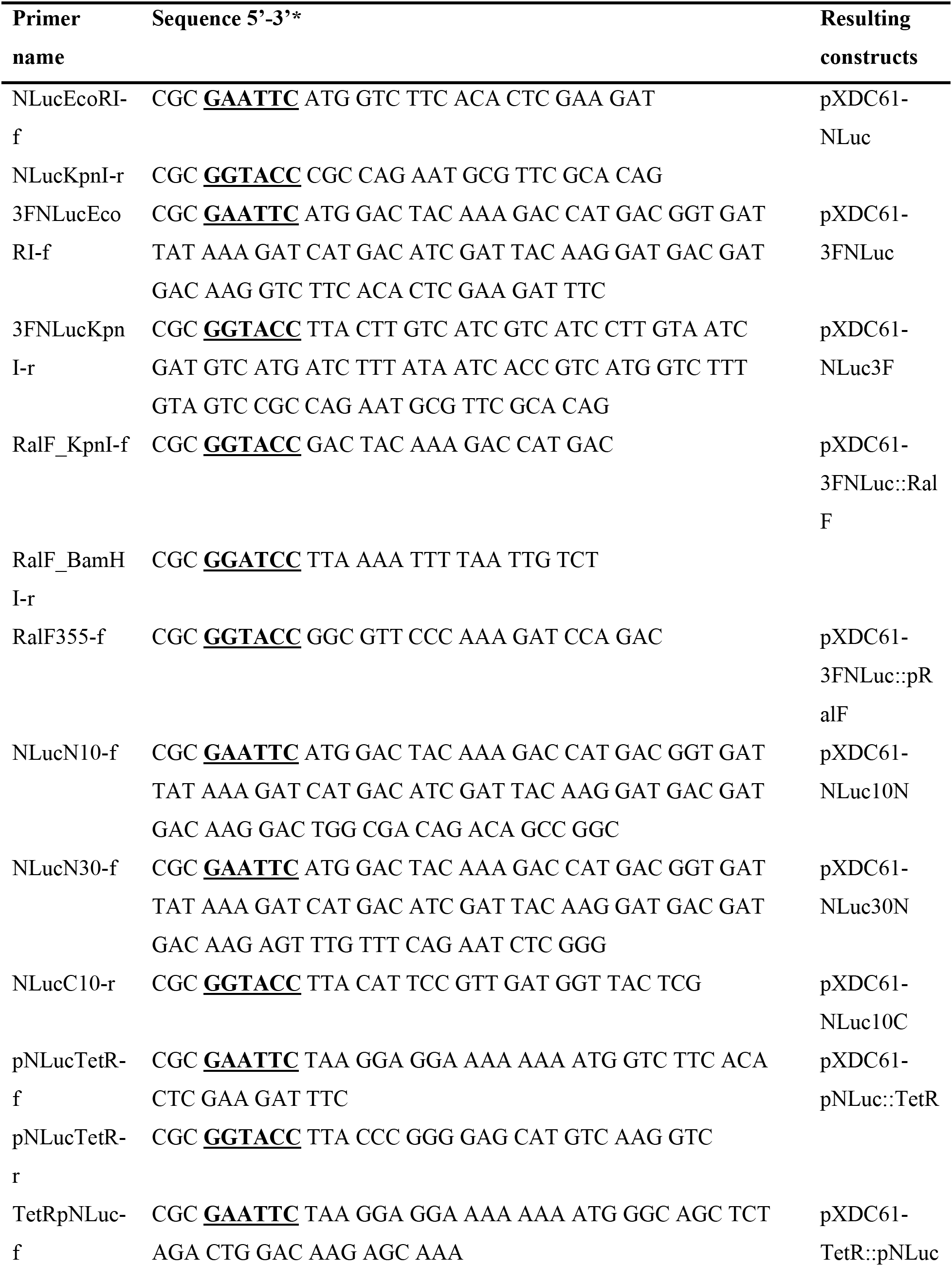
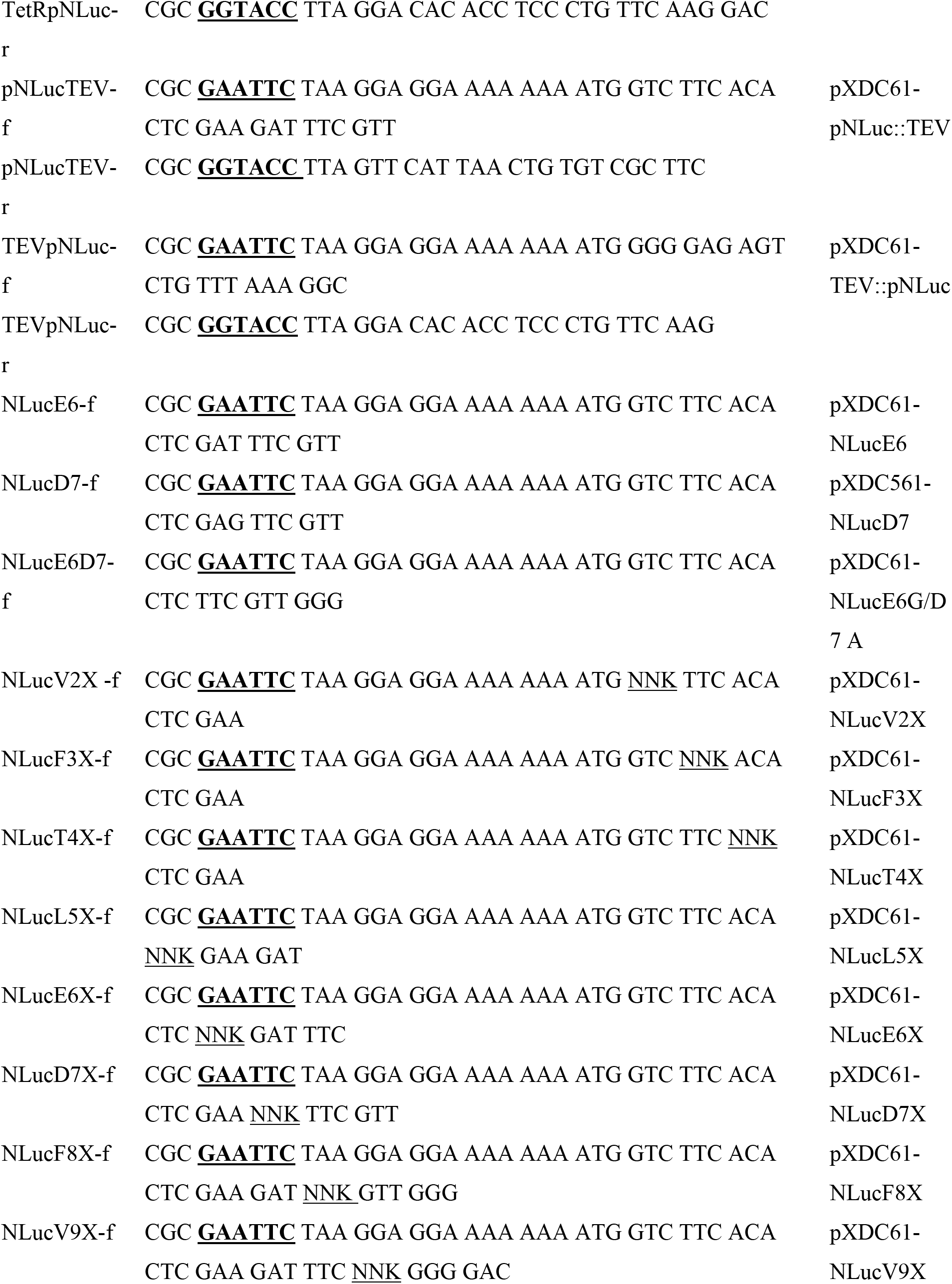
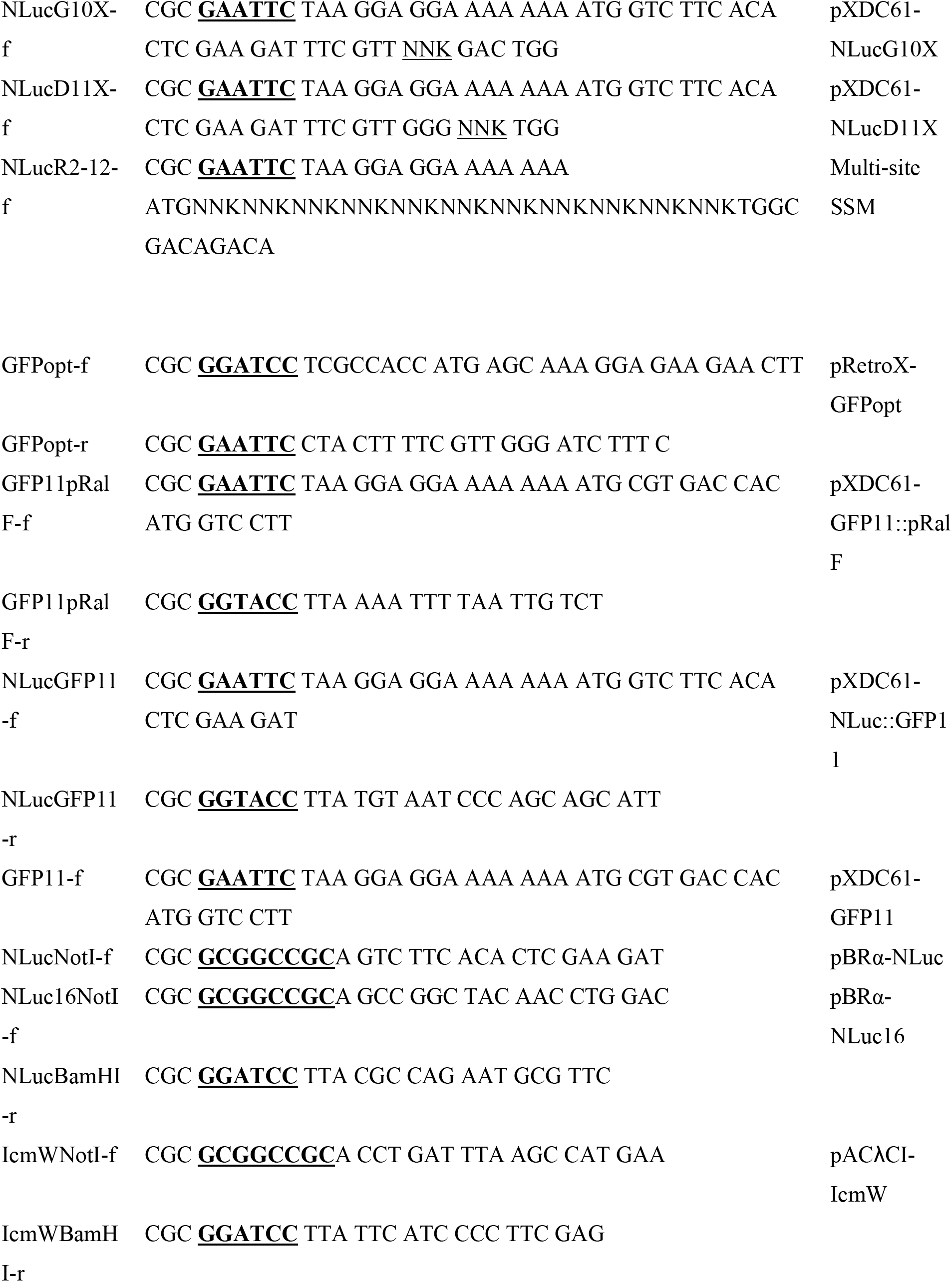
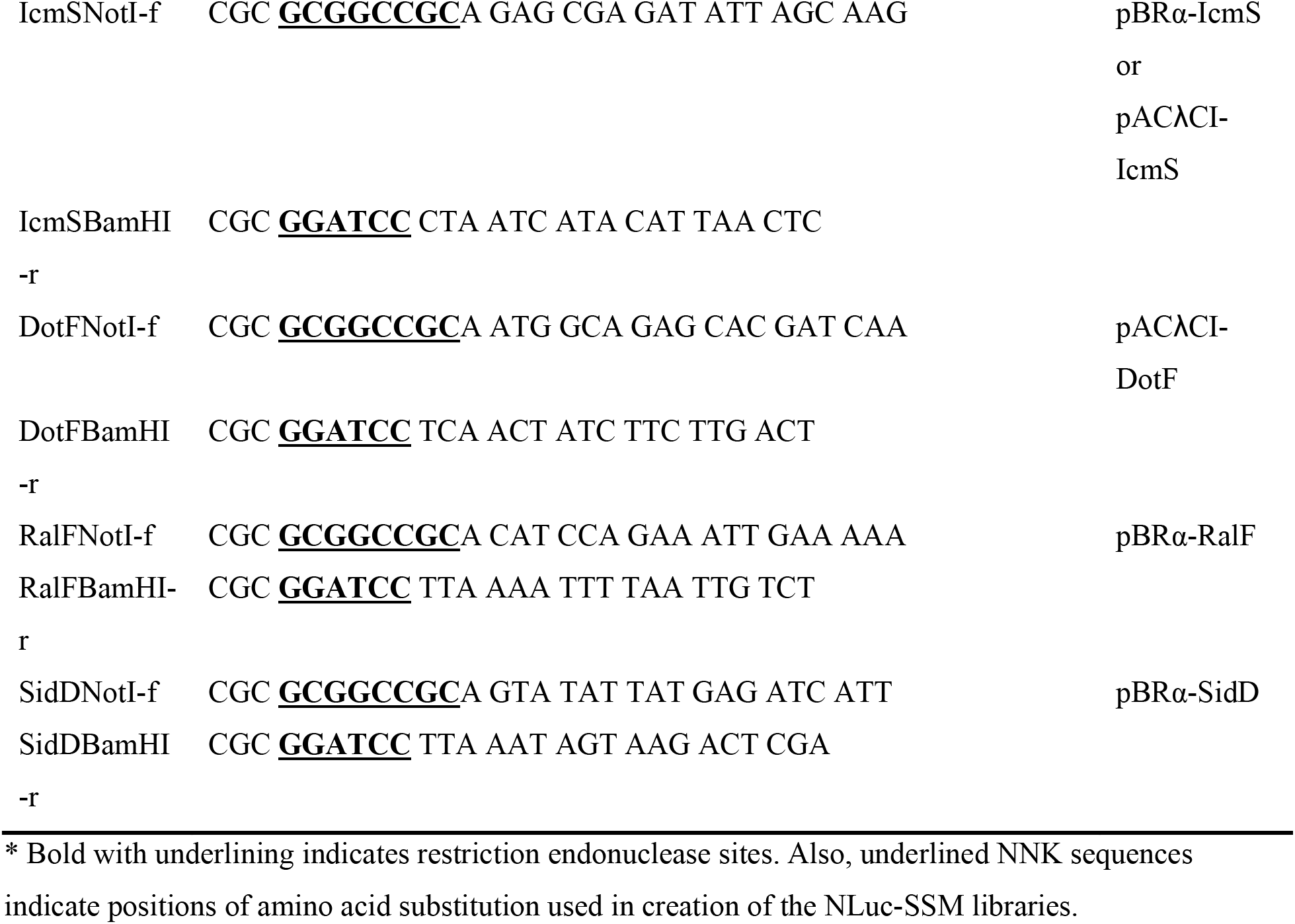
Primers used in this study.

## References

1. Christie PJ, Atmakuri K, Krishnamoorthy V, Jakubowski S, Cascales E. 2005. Biogenesis, architecture, and function of bacterial type IV secretion systems. Annu Rev Microbiol 59:451–85.

2. Lawley TD, Klimke WA, Gubbins MJ, Frost LS. 2003. F factor conjugation is a true type IV secretion system. FEMS Microbiol Lett 224:1–15.

3. Gonzalez-Rivera C, Bhatty M, Christie PJ. 2016. Mechanism and Function of Type IV Secretion During Infection of the Human Host. Microbiol Spectr 4.

4. Voth DE, Broederdorf LJ, Graham JG. 2012. Bacterial Type IV secretion systems: versatile virulence machines. Future Microbiol 7:241–57.

5. Shrivastava R, Miller JF. 2009. Virulence factor secretion and translocation by Bordetella species. Curr Opin Microbiol 12:88–93.

6. Siamer S, Dehio C. 2015. New insights into the role of Bartonella effector proteins in pathogenesis. Curr Opin Microbiol 23:80–5.

7. Ke Y, Wang Y, Li W, Chen Z. 2015. Type IV secretion system of Brucella spp. and its effectors. Front Cell Infect Microbiol 5:72.

8. Christie PJ. 2004. Type IV secretion: the Agrobacterium VirB/D4 and related conjugation systems. Biochim Biophys Acta 1694:219–34.

9. Voth DE, Heinzen RA. 2009. Coxiella type IV secretion and cellular microbiology. Curr Opin Microbiol 12:74–80.

10. Yuan XY, Wang Y, Wang MY. 2018. The type IV secretion system in Helicobacter pylori. Future Microbiol 13:1041–1054.

11. Bardill JP, Miller JL, Vogel JP. 2005. IcmS-dependent translocation of SdeA into macrophages by the Legionella pneumophila type IV secretion system. Mol Microbiol 56:90–103.

12. Sutherland MC, Nguyen TL, Tseng V, Vogel JP. 2012. The Legionella IcmSW complex directly interacts with DotL to mediate translocation of adaptor-dependent substrates. PLoS Pathog 8:e1002910.

13. Nagai H, Cambronne ED, Kagan JC, Amor JC, Kahn RA, Roy CR. 2005. A C-terminal translocation signal required for Dot/Icm-dependent delivery of the Legionella RalF protein to host cells. Proc Natl Acad Sci U S A 102:826–31.

14. Casper-Lindley C, Dahlbeck D, Clark ET, Staskawicz BJ. 2002. Direct biochemical evidence for type III secretion-dependent translocation of the AvrBs2 effector protein into plant cells. Proc Natl Acad Sci U S A 99:8336–41.

15. Sory MP, Cornelis GR. 1994. Translocation of a hybrid YopE-adenylate cyclase from Yersinia enterocolitica into HeLa cells. Mol Microbiol 14:583–94.

16. Inouye S, Watanabe K, Nakamura H, Shimomura O. 2000. Secretional luciferase of the luminous shrimp Oplophorus gracilirostris: cDNA cloning of a novel imidazopyrazinone luciferase(1). FEBS Lett 481:19–25.

17. Hall MP, Unch J, Binkowski BF, Valley MP, Butler BL, Wood MG, Otto P, Zimmerman K, Vidugiris G, Machleidt T, Robers MB, Benink HA, Eggers CT, Slater MR, Meisenheimer PL, Klaubert DH, Fan F, Encell LP, Wood KV. 2012. Engineered luciferase reporter from a deep sea shrimp utilizing a novel imidazopyrazinone substrate. ACS Chem Biol 7:1848–57.

18. England CG, Ehlerding EB, Cai W. 2016. NanoLuc: A Small Luciferase Is Brightening Up the Field of Bioluminescence. Bioconjug Chem 27:1175–1187.

19. Boute N, Lowe P, Berger S, Malissard M, Robert A, Tesar M. 2016. NanoLuc Luciferase – A Multifunctional Tool for High Throughput Antibody Screening. Front Pharmacol 7:27.

20. Thorne N, Inglese J, Auld DS. 2010. Illuminating insights into firefly luciferase and other bioluminescent reporters used in chemical biology. Chem Biol 17:646–57.

21. Meir A, Chetrit D, Liu L, Roy CR, Waksman G. 2018. Legionella DotM structure reveals a role in effector recruiting to the Type 4B secretion system. Nat Commun 9:507.

22. Christie PJ. 2017. Structural biology: Loading T4SS substrates. Nature Microbiology 2:17125.

23. Meir A, Macé K, Lukoyanova N, Chetrit D, Hospenthal MK, Redzej A, Roy C, Waksman G. 2020. Mechanism of effector capture and delivery by the type IV secretion system from Legionella pneumophila. Nat Commun 11:2864.

24. Ninio S, Zuckman-Cholon DM, Cambronne ED, Roy CR. 2005. The Legionella IcmS-IcmW protein complex is important for Dot/Icm-mediated protein translocation. Mol Microbiol 55:912–26.

25. Cambronne ED, Roy CR. 2007. The Legionella pneumophila IcmSW complex interacts with multiple Dot/Icm effectors to facilitate type IV translocation. PLoS Pathog 3:e188.

26. Heller DM, Tavag M, Hochschild A. 2017. CbtA toxin of Escherichia coli inhibits cell division and cell elongation via direct and independent interactions with FtsZ and MreB. PLoS Genet 13:e1007007.

27. Dove SL, Joung JK, Hochschild A. 1997. Activation of prokaryotic transcription through arbitrary protein-protein contacts. Nature 386:627–30.

28. Sutherland MC, Binder KA, Cualing PY, Vogel JP. 2013. Reassessing the role of DotF in the Legionella pneumophila type IV secretion system. PLoS One 8:e65529.

29. Kim H, Kubori T, Yamazaki K, Kwak MJ, Park SY, Nagai H, Vogel JP, Oh BH. 2020. Structural basis for effector protein recognition by the Dot/Icm Type IVB coupling protein complex. Nat Commun 11:2623.

30. Xu J, Xu D, Wan M, Yin L, Wang X, Wu L, Liu Y, Liu X, Zhou Y, Zhu Y. 2017. Structural insights into the roles of the IcmS–IcmW complex in the type IVb secretion system of *Legionella pneumophila*. Proc Natl Acad Sci U S A 114:13543–13548.

31. Burstein D, Zusman T, Degtyar E, Viner R, Segal G, Pupko T. 2009. Genome-scale identification of Legionella pneumophila effectors using a machine learning approach. PLoS Pathog 5:e1000508.

32. Lifshitz Z, Burstein D, Peeri M, Zusman T, Schwartz K, Shuman HA, Pupko T, Segal G. 2013. Computational modeling and experimental validation of the Legionella and Coxiella virulence-related type-IVB secretion signal. Proc Natl Acad Sci U S A 110:E707–15.

33. Huang L, Boyd D, Amyot WM, Hempstead AD, Luo ZQ, O’Connor TJ, Chen C, Machner M, Montminy T, Isberg RR. 2011. The E Block motif is associated with Legionella pneumophila translocated substrates. Cell Microbiol 13:227–45.

34. Källberg M, Wang H, Wang S, Peng J, Wang Z, Lu H, Xu J. 2012. Template-based protein structure modeling using the RaptorX web server. Nat Protoc 7:1511–22.

35. Lovell S, Mehzabeen N, Battaile KP, Wood MG, Encell LP, Wood KV. 2016. 5IBO: 1.95A resolution structure of NanoLuc luciferase. https://www.rcsb.org/structure/5ibo. Accessed March 3, 2022.

36. Mirdita M, Schütze K, Moriwaki Y, Heo L, Ovchinnikov S, Steinegger M. 2022. ColabFold – Making protein folding accessible to all. bioRxiv doi:10.1101/2021.08.15.456425:2021.08.15.456425.

37. Pédelacq JD, Cabantous S, Tran T, Terwilliger TC, Waldo GS. 2006. Engineering and characterization of a superfolder green fluorescent protein. Nat Biotechnol 24:79–88.

38. Cabantous S, Terwilliger TC, Waldo GS. 2005. Protein tagging and detection with engineered self-assembling fragments of green fluorescent protein. Nat Biotechnol 23:102–7.

39. de Felipe KS, Glover RT, Charpentier X, Anderson OR, Reyes M, Pericone CD, Shuman HA. 2008. Legionella eukaryotic-like type IV substrates interfere with organelle trafficking. PLoS Pathog 4:e1000117.

40. de Felipe KS, Pampou S, Jovanovic OS, Pericone CD, Ye SF, Kalachikov S, Shuman HA. 2005. Evidence for acquisition of Legionella type IV secretion substrates via interdomain horizontal gene transfer. J Bacteriol 187:7716–26.

41. Chen J, de Felipe KS, Clarke M, Lu H, Anderson OR, Segal G, Shuman HA. 2004. Legionella effectors that promote nonlytic release from protozoa. Science 303:1358–61.

42. Zhu W, Banga S, Tan Y, Zheng C, Stephenson R, Gately J, Luo ZQ. 2011. Comprehensive identification of protein substrates of the Dot/Icm type IV transporter of Legionella pneumophila. PLoS One 6:e17638.

43. Shohdy N, Efe JA, Emr SD, Shuman HA. 2005. Pathogen effector protein screening in yeast identifies Legionella factors that interfere with membrane trafficking. Proc Natl Acad Sci U S A 102:4866–71.

44. Luo ZQ, Isberg RR. 2004. Multiple substrates of the Legionella pneumophila Dot/Icm system identified by interbacterial protein transfer. Proc Natl Acad Sci U S A 101:841–6.

45. Lovell S, Mehzabeen N, Battaile KP, Wood MG, Encell LP, Wood KV. 1.95A resolution structure of NanoLuc luciferase. http//:10.2210/pdb5IBO/pdb. http://www.rcsb.org/structure/5IBO. Accessed

46. Buel GR, Walters KJ. 2022. Can AlphaFold2 predict the impact of missense mutations on structure? Nat Struct Mol Biol 29:1–2.

47. Chiaraviglio L, Kirby JE. 2015. High-Throughput Intracellular Antimicrobial Susceptibility Testing of Legionella pneumophila. Antimicrob Agents Chemother 59:7517–29.

48. O’Boyle N, Connolly JPR, Roe AJ. 2018. Tracking elusive cargo: Illuminating spatio-temporal Type 3 effector protein dynamics using reporters. Cell Microbiol 20.

49. Park E, Lee HY, Woo J, Choi D, Dinesh-Kumar SP. 2017. Spatiotemporal Monitoring of Pseudomonas syringae Effectors via Type III Secretion Using Split Fluorescent Protein Fragments. Plant Cell 29:1571–1584.

50. Sakalis PA, van Heusden GP, Hooykaas PJ. 2014. Visualization of VirE2 protein translocation by the Agrobacterium type IV secretion system into host cells. Microbiologyopen 3:104–17.

51. Coers J, Vance RE, Fontana MF, Dietrich WF. 2007. Restriction of Legionella pneumophila growth in macrophages requires the concerted action of cytokine and Naip5/Ipaf signalling pathways. Cell Microbiol 9:2344–57.

52. Kang YS, Kirby JE. 2017. Promotion and Rescue of Intracellular Brucella neotomae Replication during Coinfection with Legionella pneumophila. Infect Immun 85.

53. Miller JF. 1972. Experiments in molecular genetics. Cold Spring Harbor Laboratory, Cold Spring Harbor, NY.

54. Berger KH, Isberg RR. 1993. Two distinct defects in intracellular growth complemented by a single genetic locus in Legionella pneumophila. Mol Microbiol 7:7–19.

55. Zuckman DM, Hung JB, Roy CR. 1999. Pore-forming activity is not sufficient for Legionella pneumophila phagosome trafficking and intracellular growth. Mol Microbiol 32:990–1001.

